# Systemic inhibition of *de novo* purine biosynthesis prevents weight gain and improves metabolic health by increasing thermogenesis and decreasing food intake

**DOI:** 10.1101/2024.10.28.620705

**Authors:** Jacob W. Myers, Woo Yong Park, Alexander M. Eddie, Abhijit B. Shinde, Praveena Prasad, Alexandria C. Murphy, Michael Z. Leonard, Julia A. Pinette, Jessica J. Rampy, Claudia Montufar, Zayedali Shaikh, Tara T. Hickman, Garrett N. Reynolds, Nathan C. Winn, Louise Lantier, Sun H. Peck, Katie C. Coate, Roland W. Stein, Nancy Carrasco, Erin S. Calipari, Melanie R. McReynolds, Elma Zaganjor

**Affiliations:** Department of Molecular Physiology and Biophysics, Vanderbilt University School of Medicine, Nashville, TN, USA; Department of Biochemistry and Molecular Biology, Huck Institutes of the Life Sciences, Pennsylvania State University, University Park, PA; Department of Pharmacology, Vanderbilt University School of Medicine, Nashville, TN, USA; Vanderbilt Center for Addiction Research, Vanderbilt University, Nashville, TN, USA; Department of Cellular & Molecular Physiology, Yale University, New Haven, CT, USA; Department of Biochemistry, Vanderbilt University School of Medicine, Nashville, TN, USA; Department of Medicine, Vanderbilt University Medical Center, Nashville, TN, USA; Vanderbilt Mouse Metabolic Phenotyping Center, Nashville, TN, USA; Department of Biomedical Engineering, Vanderbilt University School of Engineering, Nashville, TN, USA; Department of Veterans Affairs, Tennessee Valley Healthcare System, Nashville, TN, USA; Division of Diabetes, Endocrinology and Metabolism, Department of Medicine, Vanderbilt University Medical Center, Nashville, TN, USA; Vanderbilt Digestive Disease Research Center, Vanderbilt University Medical Center, Nashville, TN, USA; Vanderbilt Diabetes Research and Training Center, Vanderbilt University Medical Center, Nashville, TN, USA

**Keywords:** thermogenesis, purine, IMPDH, mizoribine, weight, metabolism

## Abstract

**Objective:** Obesity is a major health concern, largely because it contributes to type 2 diabetes mellitus (T2DM), cardiovascular disease, and various malignancies. Increase in circulating amino acids and lipids, in part due to adipose dysfunction, have been shown to drive obesity-mediated diseases. Similarly, elevated purines and uric acid, a degradation product of purine metabolism, are found in the bloodstream and in adipose tissue. These metabolic changes are correlated with metabolic syndrome, but little is known about the physiological effects of targeting purine biosynthesis.

**Methods:** To determine the effects of purine biosynthesis on organismal health we treated mice with mizoribine, an inhibitor of inosine monophosphate dehydrogenase 1 and 2 (IMPDH1/2), key enzymes in this pathway. Mice were fed either a low-fat (LFD; 13.5% kcal from fat) or a high-fat (HFD; 60% kcal from fat) diet for 30 days during drug or vehicle treatment. We ascertained the effects of mizoribine on weight gain, body composition, food intake and absorption, energy expenditure, and overall metabolic health.

**Results:** Mizoribine treatment prevented mice on a HFD from gaining weight, but had no effect on mice on a LFD. Body composition analysis demonstrated that mizoribine significantly reduced fat mass but did not affect lean mass. Although mizoribine had no effect on lipid absorption, food intake was reduced. Furthermore, mizoribine treatment induced adaptive thermogenesis in skeletal muscle by upregulating sarcolipin, a regulator of muscle thermogenesis. While mizoribine-treated mice exhibited less adipose tissue than controls, we did not observe lipotoxicity. Rather, mizoribine-treated mice displayed improved glucose tolerance and reduced ectopic lipid accumulation.

**Conclusions:** Inhibiting purine biosynthesis prevents mice on a HFD from gaining weight, and improves their metabolic health, to a significant degree. We also demonstrated that the purine biosynthesis pathway plays a previously unknown role in skeletal muscle thermogenesis. A deeper mechanistic understanding of how purine biosynthesis promotes thermogenesis and decreases food intake may pave the way to new anti-obesity therapies. Crucially, given that many purine inhibitors have been FDA-approved for use in treating various conditions, our results indicate that they may benefit overweight or obese patients.

**Highlights:**

- A purine biosynthesis inhibitor, mizoribine, protects against diet-induced weight gain
- Mizoribine prevents fat mass gain in high-fat diet-fed male mice
- Mizoribine reduces food intake and increases thermogenesis
- Mizoribine induces expression of sarcolipin, a regulator of thermogenesis
- Mizoribine treatment reduces ectopic lipids and increases glucose tolerance

## 1. Introduction

Obesity is a chronic metabolic disease affecting 4 in 10 adults in the United States and 1 in 10 adults globally [1–3]. It is thus a substantial public health concern, as obesity is a significant risk factor for several diseases including T2DM, atherosclerotic cardiovascular disease (ACD), metabolic-associated fatty liver disease (MAFLD), and numerous cancers [2–6]. A variety of treatments exist for managing obesity including lifestyle changes, bariatric surgeries, and pharmacological interventions. Anti-obesity medications are typically considered when lifestyle changes alone do not enable a patient to achieve and maintain weight loss, and when the health risks associated with obesity are significant [7]. These medications work through a variety of mechanisms, including suppressing appetite, altering nutrient absorption, and affecting the hormonal regulation of body weight. Such medications are, however, of limited efficacy, and they often have substantial adverse effects [8–10]. Thus, obesity and obesity-mediated diseases remain prevalent health problems.

Obesity comes about when white adipose tissue expands in response to positive energy balance. This imbalance occurs when energy intake exceeds energy expenditure, causing surplus energy to be stored as fat. Adipose tissue serves a protective function by safely storing excess nutrients and preventing lipotoxicity. However, in the course of chronic overnutrition, adipose tissue itself becomes dysfunctional. It is characterized by enlarged adipocytes, hypoxia, inflammation, and fibrosis [11–14]. Collectively, these changes in adipose tissue contribute to systemic metabolic dysfunction. Perhaps most notably, dysfunctional adipose tissue exhibits increased lipolysis which leads to elevated plasma levels of free fatty acids, ectopic lipid accumulation in the liver, and reduced insulin sensitivity [15]. In addition, numerous laboratories have found that increased blood levels of branched-chain amino acids (BCAAs) and aromatic amino acid concentrations are highly predictive of patients’ risk of developing T2DM and that reducing BCAA levels improves metabolic health [16,17]. Taken together, these studies strongly suggest that it is useful to identify and target metabolites that are systemically altered in obesity.

Recent developments in systems epidemiology have led to the discovery of changes in circulating and adipose tissue metabolites that correlate with obesity and obesity-mediated diseases [15]. These include changes in byproducts of purine metabolism, such as uric acid, GMP, GTP, IMP, guanosine, xanthosine, and inosine [18–21]. Reduction in visceral fat during weight loss is associated with reduced uric acid levels suggesting that white adipose tissue may be an important site of purine metabolism [22]. Furthermore, hyperuricemia induces oxidative stress and upregulation of proinflammatory cytokines in adipose tissue and may be conducive to insulin resistance [20,23–26]. Increases in purine levels may be due to increased uptake from food, from *de novo* purine biosynthesis, or from the nucleotide salvage pathway. Because specific pharmacological approaches to inhibit *de novo* purine biosynthesis are well established, we investigated how perturbations in this pathway affect diet-induced obesity and the associated metabolic dysfunction. We found that treating mice with daily injections of a purine biosynthesis inhibitor, mizoribine, prevents them from gaining weight even when they are on a HFD - — and, in particular, decreases their fat mass accumulation. Our results further establish that systemically blocking *de novo* nucleotide biosynthesis prevents HFD-mediated weight gain by decreasing food intake and increasing thermogenesis. Although disrupting adipose tissue can induce lipotoxicity in other organs, we find that mizoribine-mediated decrease in fat mass is not associated with increased ectopic lipid accumulation. Rather, mizoribine treatment improves glucose tolerance in HFD-fed mice suggesting that it has a positive effect on the metabolic health of mice facing a dietary challenge. Taken together, our results reveal a previously unknown physiological effect of purine synthesis inhibition: it protects against HFD-mediated dysfunction.

## 2. Methods

### 2.1 Diet Challenge & Mizoribine Treatment

For all studies, age-matched, wild type, C57BL/6J mice were used. Mice were between 8 and 12 weeks of age at the start of experimental manipulation. Unless otherwise noted, mice were treated for 30 days with daily intraperitoneal injections of either 50 milligrams per kilogram body weight mizoribine (MedChemExpress; catalog no.: HY-17470) in 2% DMSO in phosphate-buffered saline (PBS) or a control vehicle injection of 2% DMSO in PBS. Mice were given water and placed on either a low-fat diet (LFD; LabDiet; catalog no.: 5LOD) or a high-fat diet (HFD; Research Diets; catalog no.: D12492) *ad libitum*. The LFD contained 13.5% energy from fat, whereas the HFD contained 60% energy from fat. Mice remained on a constant 12-hour light, 12-hour dark cycle throughout the experiment and were housed with 3-5 animals in a cage. Mouse body weights and food consumption were recorded daily. Body temperature was measured daily by infrared thermometry. All experimental protocols were approved by the Vanderbilt University Institutional Animal Care and Use Committee as required by the Public Health Service Policy on Humane Care and Use of Laboratory Animals.

### 2.2 Dual Energy X-ray Absorptiometry

On day 0 and day 30, the body composition of each mouse was assessed using dual-energy x-ray absorptiometry (DXA). Each mouse was anesthetized for the duration of the procedure using 2% isoflurane delivered via nose cone. Mice were placed in the DXA scanner in a prone position, and imaged using a Faxitron UltraFocus (Hologic, Inc. Marlborough, MA). The instrument was calibrated for offset images, dark images, and flat-field images before the measurement using a method provided by the manufacturer. For the purpose of assessing body composition, the cranium was excluded from measurements. Measures of lean weight, fat weight, fat percentage, and lean percentage were calculated from segmented regions of interest (ROIs) in the Vision DXA software (Version 2.4.2U). Pseudo-color visualizations of bone, lean, and fat mass distribution were obtained by overlaying radiopacity maps of these ROIs generated in Vision DXA using the Merge Channels tool in ImageJ (Version 1.54f) [27].

### 2.3 Tissue Collection

At the conclusion of the experiment (Day 30), mice were euthanized by isoflurane inhalation and cervical dislocation. Immediately after euthanasia, intracardiac blood was collected. Plasma was isolated by adding blood to a tube containing 0.1 M Sodium ethylenediaminetetraacetic acid (10% of total sample volume) and centrifuging at 20000 x *g* for 20 minutes. The left and right perigonadal visceral white adipose tissue (VAT), inguinal subcutaneous white adipose tissue (SAT), and interscapular brown adipose tissue (BAT) depots, as well as the liver, pancreas, quadriceps (QUAD), soleus (SOL), gastrocnemius (GAS), and tibialis anterior (TA) were isolated. These tissues were weighed, and the inguinal lymph node was removed from the SAT depot. Portions of each tissue were subsequently fixed in 10% neutral buffered formalin for a minimum of 48 hours before being embedded in paraffin. Portions of the liver were frozen in cryomolds in optimal cutting temperature tissue freezing medium (Sakura; Catalog no.: 4583). Additional portions of these tissues were snap frozen in liquid nitrogen, powderized, and stored at –80 °C prior to immunoblot and RT-qPCR analyses.

### 2.4 Histology & Microscopy

Formalin-fixed, paraffin-embedded tissue blocks were serially cut to 5 µm sections and stained with hematoxylin and eosin (H&E). Images of the H&E-stained tissue sections were acquired using an Aperio ScanScope CS brightfield digital scanner (Aperio ScanScope CS, Aperio Technologies, Leica Biosystems, Deer Park, IL) at 20x magnification.

### 2.5 Image Analysis

To identify morphological changes in white adipose tissue, average adipocyte size was measured from images of H&E-stained tissue sections. Images were segmented using QuPath’s Pixel Classifier tool (Version 0.4.3) using the random trees method at full resolution and with all available features and scales [28]. Segmented adipocytes were then quantified using ImageJ’s Analyze Particles tool (Version 1.54f) with settings size = 250 – 50000 µm^2^ and circularity = 0.1 – 1.0 [27].

To identify morphological changes in brown adipose tissue and the liver, average lipid droplet size was measured from images of H&E stained tissue sections using ImageJ (Version 1.54f) [27]. Color deconvolution was first applied to isolate the eosin signal. Appropriate thresholding was applied to the eosin signal to segment the lipid droplets and the images were subsequently binarized. Additionally, for the brown adipose tissue, watershed segmentation was performed on the binary images. Segmented lipid droplets from both liver and brown adipose tissue were measured using the Analyze Particles tool.

### 2.6 Fecal Lipid Composition

On day 30, mice were placed in individual housing for 2 hours and feces were collected. Feces were frozen at -80° C and powderized. Free fatty acids (FFAs) were extracted from powderized samples as previously described [29]. The extracts were filtered and lipids were recovered from the chloroform phase. Individual lipid classes were separated by thin-layer chromatography using Silica Gel 60 A plates developed in petroleum ether, ethyl ether, and acetic acid (80:20:1) and visualized by rhodamine 6G. Free fatty acids were scraped from the plate and methylated using BF_3_ /methanol as previously described [30]. The fatty acid methyl esters were analyzed by gas chromatography using an Agilent 7890 gas chromatograph equipped with a flame ionization detector and a capillary column (SP2380, 0.25 mm x 30 m, 0.20 µm film, Supelco, Bellefonte, PA). Helium was used as a carrier gas, and the oven temperature was programmed from 160 °C to 230 °C at 4 °C/min. Fatty acid methyl esters were identified by comparing the retention times to those of known standards, and inclusion of a pentadecanoic acid (15:0) internal standard permitted quantitation of the amount of FFA in the sample.

### 2.7 Enzyme-linked Immunosorbent Assay for Leptin

Plasma leptin concentrations were measured using the Murine Leptin Standard ABTS ELISA Development Kit (Peprotech 900-K76). Methods were followed per the manufacturer’s instructions. Samples and standards were incubated overnight at 4 °C. All samples were diluted 1:100 in 2% bovine serum albumin (BSA) in PBS and plated in duplicate. Absorbance was detected at 405 nm using the ThermoScientific Varioscan LUX. Sample concentrations were calculated by generating a standard curve from known recombinant leptin concentrations.

### 2.8 Conditioned Taste Aversion

Mice were habituated to the experimental environment and handling procedures for a minimum of 3 days prior to the start of the experiment. Mice were individually housed and given *ad libitum* access to water and LFD throughout the experiment. For the first 7 days of the experiment (Days 0 – 6), mice were exposed to a novel, flavored liquid (Flavor A; Abbott; Catalog no.: 53432) for a period of 2 hours. For the next 7 days of the experiment (Days 7 – 13), mice were intraperitoneally injected with either 50 mg per kg body weight mizoribine in 2% DMSO in PBS or a control vehicle injection of 2% DMSO in PBS. Immediately following injection, mice were exposed to a second novel, flavored liquid (Flavor B; Abbott; Catalog no.: 53623) for a period of 2 hours. After 7 days of injections and flavor B exposure (Day 14), mice were exposed to Flavor A for 2 hours and consumption was measured as a control. On the final day of the experiment (Day 15), mice were exposed to Flavor B for 2 hours and consumption was measured. Taste aversion was assessed by comparing the consumption of Flavor B on Day 15 between the mizoribine and vehicle treatment groups.

### 2.9 Indirect Calorimetry & Locomotor Activity

To identify changes in energy balance, food intake, and energy expenditure were measured using a Promethion Core Metabolic System (Sable Systems International, Las Vegas, NV). The system is housed in a temperature and light-controlled cabinet (temperature maintained at 22 °C during the run) in a dedicated room. In one experiment, male mice were transferred to the Promethion system for 7 days. While in the Promethion system, these mice were challenged with a HFD and treated with daily injections of mizoribine or vehicle. In a second experiment, male mice were challenged with a HFD and treated with daily injections of mizoribine or vehicle for 30 days as previously described. On day 30, mice were moved to the Promethion system for 7 days. Additionally, body temperature was monitored using implanted temperature probes (IPTT-300, Bio Medic Data Systems). While in the Promethion system, mice continued to be challenged with a HFD and received daily injections of mizoribine or vehicle. Temperature was measured daily, beginning on day 34, at exactly 8 hours into the light photoperiod. In both experiments, O_2_ consumption, CO_2_ production, water consumption, and locomotion were continually measured. In both experiments, mice were given 3 days to acclimate before indirect calorimetry measurements were taken. The respiratory exchange ratio (RER) was calculated by dividing the CO_2_ production rate by the O_2_ consumption rate. Energy expenditure was calculated from the O_2_ consumption and CO_2_ production rates using the Weir Equation [31].

### 2.10 Acute Cold Exposure Challenge

Mice were maintained at 22 °C throughout the 30 day diet and drug treatment period. After 30 days of treatment, body temperature was measured using infrared thermometry. Mice were then placed at 4 °C and body temperature was measured hourly for six hours.

### 2.11 Chemiluminescent Immunoassays for Thyroid Hormones

Plasma was isolated from intracardiac blood by centrifugation. Samples were diluted 1:10 in 1x PBS and the concentration of thyroid hormones T_3_ and T_4_ was measured from the plasma using chemiluminescent immunoassays (Diagnostic Automation; Catalog no.: 9003-16 and 9001-16) according to the manufacturer’s protocol.

### 2.12 Immunoblotting

Frozen, powderized tissues samples were lysed with radioimmunoprecipitation assay lysis buffer (1% NP-40, 150 mM NaCl, 25 mM Tris base, 0.5% sodium deoxycholate, 0.1% SDS, 1% phosphatase inhibitor cocktails #2 and #3 (Sigma-Aldrich; Catalog no.: P0044 and P5726), one cOmplete protease inhibitor tablet (Sigma-Aldrich; Catalog no.: 4693116001)). Protein content was quantified using a Bicinchoninic Acid assay (Thermo Scientific; Catalog no.: 23227), and equal protein was run on 4 to 20% Tris–Glycine Gels (Invitrogen; Catalog no.: WXP42012BOX) with a 1x SDS running buffer (0.1% SDS, 25 mM Tris, 192 mM glycine) for 140 minutes at 110 V. Protein was transferred to a nitrocellulose membrane (LICOR Biosciences; Catalog no.: 92631092) with a 1x Tris-Glycine transfer buffer (25 mM Tris, 192 mM glycine, 20% methanol) over 40 minutes at 25 V. Membranes were incubated with the following primary antibodies overnight at 4 °C: UCP1 (CST; catalog no.: 72298S), p-HSL Ser660 (CST; catalog no.: 45804), HSL (CST; catalog no.: 4107T), p-ACC Ser79 (CST; catalog no.: 3661), ACC (CST; catalog no.: 3662S), CPT1α (Invitrogen; catalog no.: PA5-106270), CKB (Abclonal; catalog no.: A12632), tubulin (Novus; catalog no: NB100-690), vinculin (Santa Cruz, catalog no.: sc-25336). Membranes were washed and incubated with secondary antibodies for one hour (Invitrogen; catalog no.: A32802 and A32789). Proteins were detected by fluorescence and quantified using Image Lab (version 6.0.0, Bio-Rad Laboratories, Inc.).

To investigate sarcolipin (SLN) expression, site-directed polyclonal antibodies against the carboxy-terminal 6 residues were generated and purified by affinity chromatography [32]. SLN antibodies were diluted 1:1500. Immunoblotting procedures were followed as described above, with the following modifications: when examining SLN in the soleus, a 10 to 20% tricine gel (Invitrogen; catalog no.: EC6625BOX) was run with a 10x tris-tricine-SDS running buffer (Bio-Rad; catalog no.: 1610744) for 90 minutes at 200 V, and protein was electrotransferred for 16 minutes at 150 mA. When examining SLN in the diaphragm, a 4 to 12% Bis-Tris gel (Invitrogen; catalog no.: NP0321BOX) was run with a 20x MES SDS running buffer (Invitrogen; catalog no.: NP0002) for 25 minutes at 200 V, and protein was transferred using a 20% methanol 20x transfer buffer (Invitrogen; catalog no.: NP00061) for 60 minutes at 10 V.

### 2.13 RNA Isolation & Gene Expression Analysis

RNA was extracted from frozen, powderized tissue using TRIzol reagent as per the manufacturer’s instructions. Complementary DNA was synthesized from 0.5 μg of RNA using the iScript complementary DNA synthesis kit (Bio-Rad; catalog no.: 1708891). Real-time quantitative PCR was performed on a Bio-Rad CFX96 using SsoAdvanced Universal SYBR Green SuperMix (Bio-Rad; catalog no.: 1725120). The following mouse PCR primers were used:

16S rRNA:

Forward: CCGCAAGGGAAAGATGAAAGAC

Reverse: TCGTTTGGTTTCGGGGTTTC

*Rps16:*

Forward: CACTGCAAACGGGGAAATGG

Reverse: CACCAGCAAATCGCTCCTTG

*Acaca*:

Forward: TGACAGACTGATCGCAGAGAAAG

Reverse: TGGAGAGCCCCACACACA

*Fasn*:

Forward: CAGCAGAGTCTACAGCTACCT

Reverse: AACACCAGAGACCGTTATGC

*Hmgcr*:

Forward: CTTGTGGAATGCCTTGTGATTG

Reverse: AGCCGAAGCAGCACATGAT

*Hmgcs1*:

Forward: GCCGTGAACTGGGTCGAA

Reverse: GCATATATAGCAATGTCTCCTGCAA

*Ppargc1a*:

Forward: CCCTGCCATTGTTAAGACC

Reverse: TGCTGCTGTTCCTGTTTTC

*Prdm16*:

Forward: CAGCACGGTGAAGCCATTC

Reverse: GCGTGCATCCGCTTGTG

*Scd1*:

Forward: GAAGTCCACGCTCGATCTCA

Reverse: TGGAGATCTCTTGGAGCATGTG

*Srebf1c*:

Forward: GGAGCCATGGATTGCACATT

Reverse: GGCCCGGGAAGTCACTGT

*Srebf1a*:

Forward: GGCCGAGATGTGCGAACT

Reverse: TTGTTGATGAGCTGGAGCATGT

*Ucp1*:

Forward: GGCCTCTACGACTCAGTCCA

Reverse: TAAGCCGGCTGAGATCTTGT

### 2.14 Intraperitoneal Glucose Tolerance Test

Male mice were challenged with a HFD and treated with daily injections of mizoribine or vehicle for 30 days as previously described. After a 15 hour fast, basal blood glucose was measured using a handheld glucometer (Fisher; catalog no.: 23-111-390). Mice were intraperitoneally injected with 2 grams of dextrose per kilogram body weight. Blood glucose was sampled via tail cut and measured at 15-, 30-, 60-, and 120-minutes post-injection. To quantify, baseline-adjusted areas under the curve were calculated using the trapezoidal rule.

### 2.15 Insulin Radioimmunoassay

Insulin concentration was analyzed from plasma collected from the tail vein during the glucose tolerance test 15 minutes after the dextrose injection. Insulin concentration was determined using a radioimmunoassay kit (Millipore; catalog no.: PI-13K). The assay utilizes I125-labled insulin and a double antibody technique to determine plasma insulin levels. Methods were followed per the manufacturer’s instructions.

### 2.16 Immunofluorescence Staining and Image Analysis

Pancreatic tissue preparation and sectioning were performed as described previously [33,34]. Tissue sections were immunostained for insulin (Dako; catalog no.: A0564), glucagon (Cell Signaling; catalog no.: 2760), and somatostatin (American Research Products, Inc.; catalog no.: 13-2366) to label beta, alpha, and delta cells, respectively. Whole pancreatic sections were imaged using an Axio Scan.Z1 (Zeiss) slide scanning system and analyzed using Imaris (Oxford Instruments). Fractional areas for alpha, beta, and delta cells from up to 3 sections of differing tissue depths per animal were determined by dividing the glucagon+, insulin+, and somatostatin+ areas by the total tissue section area.

### 2.17 Neutral Lipid Staining and Microscopy

Neutral lipids were detected in the liver by BODIPY 493/503 (Cayman; catalog no.: 25892). Frozen liver blocks were serially cut to 5 µm sections and washed 3 times with PBS. Tissues were incubated with a 1:1000 diluted BODIPY solution for 40 minutes. The tissues were then washed 3 times with PBS. Coverslips were then mounted to slides using ProLong Gold with DAPI (Invitrogen; catalog no: P36931) and allowed to cure for 24 hours prior to imaging. Images were acquired using a Zeiss LSM 710 confocal microscope at 40x magnification.

### 2.18 Lipidomic Profiling

Detailed methods on plasma lipid extraction, sample reconstitution, LC-MS experiments, and lipidomic analysis are described below.

#### 2.18.1 Plasma extraction for lipids

For lipidomic profiling, plasma samples from male mice treated with a HFD and vehicle or mizoribine injections were used. Plasma was mixed with 1 mL of Extraction Buffer containing IPA/H2O/Ethyl Acetate (30:10:60, v/v/v) and Avanti Lipidomix Internal Standard (diluted 1:1000) (Avanti Polar Lipids, Inc. Alabaster, AL). Samples were vortexed and transferred to bead mill tubes for homogenization using a VWR Bead Mill at 6000 g for 30 seconds, repeated twice. The samples were then sonicated for 5 minutes and centrifuged at 15,000 g for 5 minutes at 4 °C. The upper phase was transferred to a new tube and kept at 4 °C. To re-extract the tissues, another 1 mL of Extraction Buffer (30:10:60, v/v/v) was added to the plasma pellet-containing tube. The samples were vortexed, homogenized, sonicated, and centrifuged as described earlier. The supernatants from both extractions were combined, and the organic phase was dried under liquid nitrogen gas.

#### 2.18.2 Sample reconstitution for lipids

The dried samples were reconstituted in 300 µL of Solvent A (IPA/ACN/H2O, 45:35:20, v/v/v). After brief vortexing, the samples were sonicated for 7 minutes and centrifuged at 15,000 x *g* for 10 minutes at 4 °C. The supernatants were centrifuged again for 5 minutes at 15,000 x *g* at 4 °C to remove any remaining particulates. For LC-MS lipidomic analysis, 60 µL of the sample extracts were transferred to mass spectrometry vials.

#### 2.18.3 LC-MS analysis for lipids

Sample analysis was performed within 36 hours after extraction using a Vanquish UHPLC system coupled with an Orbitrap Exploris 240™ mass spectrometer equipped with a H-ESI™ ion source (all Thermo Fisher Scientific). A Waters (Milford, MA) CSH C18 column (1.0 × 150 mm × 1.7 µm particle size) was used. Solvent A consisted of ACN:H2O (60:40; v/v) with 10 mM Ammonium formate and 0.1% formic acid, while solvent B contained IPA:ACN (95:5; v/v) with 10 mM Ammonium formate and 0.1% formic acid. The mobile phase flow rate was set at 0.11 mL/min, and the column temperature was maintained at 65 °C. The gradient for solvent B was as follows: 0 min 15% (B), 0–2 min 30% (B), 2–2.5 min 48% (B), 2.5–11 min 82% (B), 11–11.01 min 99% (B), 11.01–12.95 min 99% (B), 12.95–13 min 15% (B), and 13–15 min 15% (B). Ion source spray voltages were set at 4,000 V and 3,000 V in positive and negative mode, respectively. Full scan mass spectrometry was conducted with a scan range from 200 to 1000 m/z, and AcquireX mode was utilized with a stepped collision energy of 30% with a 5% spread for fragment ion MS/MS scan.

#### 2.18.4 Lipidomic analysis

Data were analyzed using R (Version 4.4.1) and the *lipidr* package (Version 2.18.0) [35]. All code to analyze data and generate figures can be found at https://github.com/jacobwm/Mizoribine-Lipidomics. For lipids with multiple transitions, a single feature was determined by selecting the maximum intensity. Probabilistic quotient normalization and log transformation were applied using the “normalize_pqn” function in *lipidr*. Principle component analysis (PCA) was performed using the “mva” function in *lipidr*. Lipid composition of samples from vehicle- and mizoribine-treated mice were compared using the “de_analysis” function in *lipidr*. The “lsea” function in *lipidr* was used to perform lipid set enrichment analysis.

### 2.19 Statistical Analysis

Descriptions of the specific statistical tests used, and the number of replicates (n) can be found for each experiment in the respective figure legend. GraphPad Prism 8 (Graphpad Software Inc, La Jolla, CA) and MS Excel were used for all statistical analyses.

## 3. Results

### 3.1 Inhibition of purine biosynthesis reduces diet-mediated fat expansion

Metabolomic studies have shown a correlation between obesity and elevation in circulating and adipose tissue purine metabolites [18–21,36]. Moreover, previous *in vitro* studies have shown that inhibition of purine biosynthesis suppresses lipid accumulation in adipocytes [37,38]. To determine whether modulating purine biosynthesis has an effect on diet-induced adiposity, we blocked inosine monophosphate dehydrogenase 1 and 2 (IMPDH1/2), critical enzymes in *de novo* purine biosynthesis, using a small molecule inhibitor, mizoribine (Fig. 1 A). We treated mice with daily intraperitoneal injections of mizoribine for 30 days. Mice were fed either a high- or a low-fat diet throughout the experiment (Fig. 1 B). Among both males and females fed a high-fat diet (HFD), mice treated with mizoribine were protected from weight gain (Fig. 1 C – F), whereas mice fed a low-fat diet (LFD) did not show significantly different changes in body weight (Fig. 1 C – F, S1 I – L). Notably, mice on a LFD treated with vehicle injections did not gain weight over the course of 30 days, possibly due to the stress of daily injections (Fig. 1 C, S1 I). Although the male mice on a LFD had a higher body mass at the beginning of the experiment than male mice on a HFD (S1 I), this is only reflective of slight differences in the age of the mice at the start of the experiment. The effects of mizoribine on male mice fed a HFD were observed across multiple cohorts; these were unaffected by slight variations in age and initial body mass at the beginning of the experiment. To examine the effect of mizoribine treatment on body composition, we performed dual-energy X-ray absorptiometry (DXA) and observed that, while treatment did not alter lean mass, fat mass was significantly reduced in male mice treated with mizoribine, independently of their diet (Fig. 1 G – L, S1 A – D, M). In female mice however, changes in body composition were more limited (Fig. 1 M – R, S1 E – H, N), likely because female mice are more resistant to fat accumulation and hyperphagia in response to short-term HFDs, as previously described [39]. These studies showed that blocking purine biosynthesis most sharply decreases the fat mass of male mice fed a HFD.

**Figure 1.**
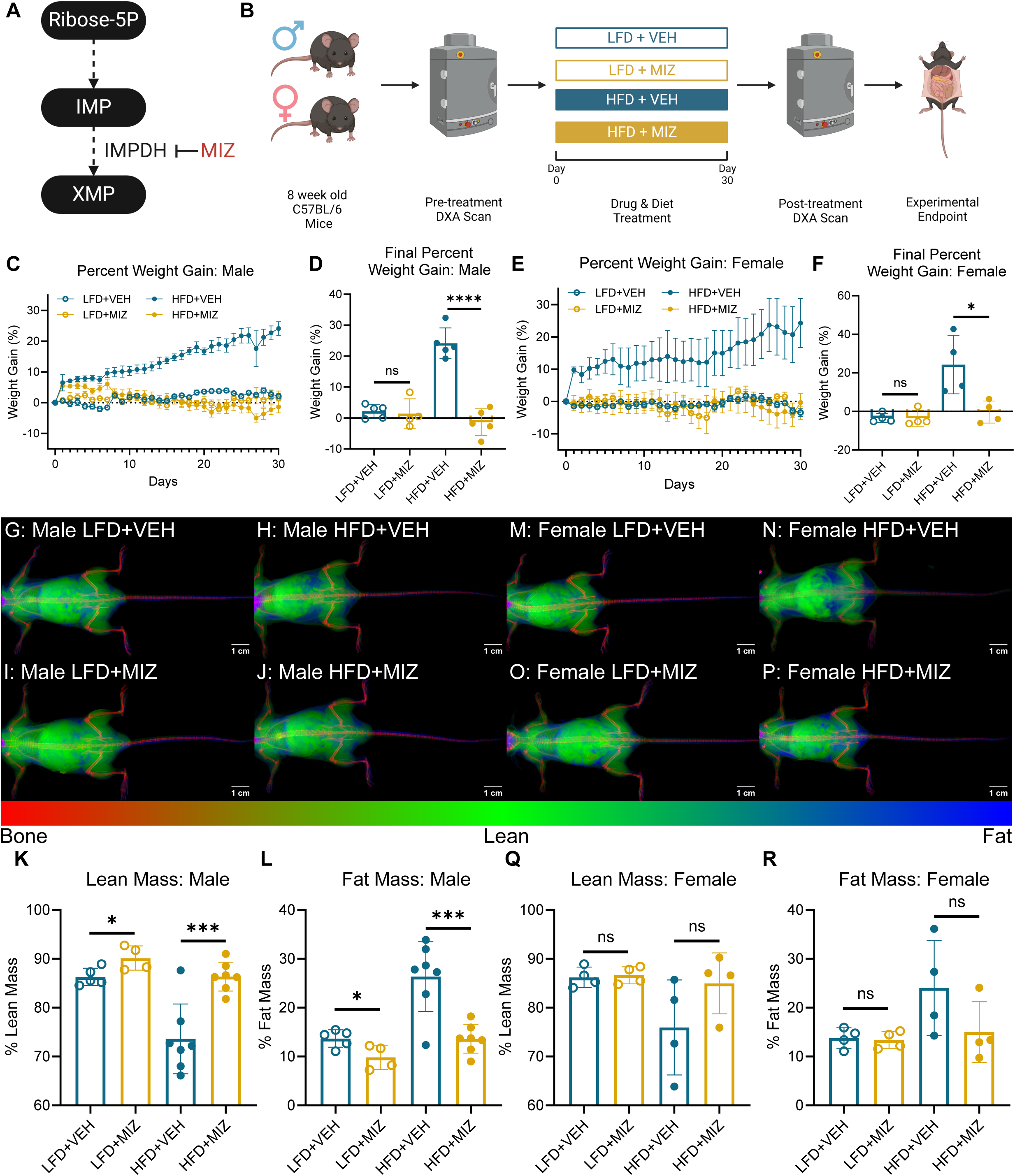
Mizoribine Treatment Disrupts Diet Mediated Fat Accumulation. (A) Abbreviated schematic of the purine biosynthesis pathway. (B) Schematic of the experimental setup. (C) Percent body weight change over 30 days in male mice fed a high fat diet (HFD) or low fat diet (LFD) and treated with daily injections of mizoribine (MIZ) or vehicle (VEH). Error bars represent mean ± standard error of the mean. N = 4-5 per group. (D) Area under the percent body weight change vs time curves in C. Error bars represent mean ± standard deviation. Significance indicative of two-tailed student’s *t*-test. Data representative of at least 3 independent experimental cohorts. (E) Percent body weight change over 30 days in female mice fed a HFD or LFD and treated with daily injections of MIZ or VEH. Error bars represent mean ± standard error of the mean. N = 4 per group. (F) Area under the percent body weight change vs time curves in E. N = 4 per group. Error bars represent mean ± standard deviation. Significance indicative of two-tailed student’s *t*-test. Data representative of at least 2 independent experimental cohorts. (G – J) Representative pseudo-color images from day 30 dual-energy X-ray absorptiometry (DXA) scans of male mice treated with LFD and vehicle (G), HFD and vehicle (H), LFD and mizoribine (I), HFD and mizoribine (J). Scale bars = 1 cm. (K – L) Quantifications of lean mass percentage (K) and fat mass percentage (L) from DXA scans in male mice. N= 4 – 7 per group. Error bars represent mean ± standard deviation. Significance indicative of two-tailed student’s *t*-test. Data representative of at least 2 independent experimental cohorts. (M – P) Representative pseudo-color images from day 30 dual-energy X-ray absorptiometry (DXA) scans of female mice treated with LFD and vehicle (M), HFD and vehicle (N), LFD and mizoribine (O), HFD and mizoribine (P). (Q – R) Quantifications of lean mass percentage (Q) and fat mass percentage (R) from DXA scans in female mice. N = 4 per group. Error bars represent mean ± standard deviation. Significance indicative of two-tailed student’s *t*-test. ns > 0.05, ∗ *p* ≤ 0.05, ∗∗ *p* ≤ 0.01, ∗∗∗ *p* ≤ 0.001, and ∗∗∗∗ *p* ≤ 0.0001.

### 3.2 Mizoribine treatment reduces adipose tissue mass and adipocyte size in male but not female mice

Considering that DXA indicated that the reduction in body weight was specific to fat mass, we examined several depots of adipose tissue. We found that male mice treated with mizoribine had significantly reduced white adipose tissue (WAT) mass in the visceral and inguinal subcutaneous depots independently of their diet (Fig. 2 A – B). Conversely, brown adipose tissue (BAT) mass was unaltered by mizoribine treatment in male mice (Fig. 2 C). We did not observe a change in WAT mass between treatment groups in female mice on either diet but found that brown adipose mass was reduced and white adipose mass, while not significantly altered, was trending down in mizoribine-treated mice on a HFD (Fig. S2 A – C). Given the established relationship between adipocyte hypertrophy and adipose dysfunction in obesity-mediated disease, we investigated white adipocyte morphology [11]. We found that white adipocytes in the subcutaneous and visceral depots were smaller in HFD-fed male mice treated with mizoribine (Fig. 2 D – F, H – I). Turning to the LFD-fed male mice, no change in visceral adipocyte size was observed in the treatment groups (Fig. 2 D – E, H); however, LFD-fed, mizoribine-treated, male mice had smaller subcutaneous white adipocytes (Fig. 2 D, F, I). By contrast, female mice—regardless of diet—showed no change in white adipocyte size, although visceral adipocyte size was trending down under mizoribine treatment when fed a HFD (Fig. S2 D – F, H – I). We also examined brown adipose tissue morphology and found that brown adipocyte lipids droplet size was unaltered by mizoribine treatment in both sexes and across all diets (Fig. 2 D, G, J; Fig. S2 D, G, J). Collectively, mizoribine most significantly reduced the white adipose depot weight and adipocyte size in male mice.

**Figure 2.**
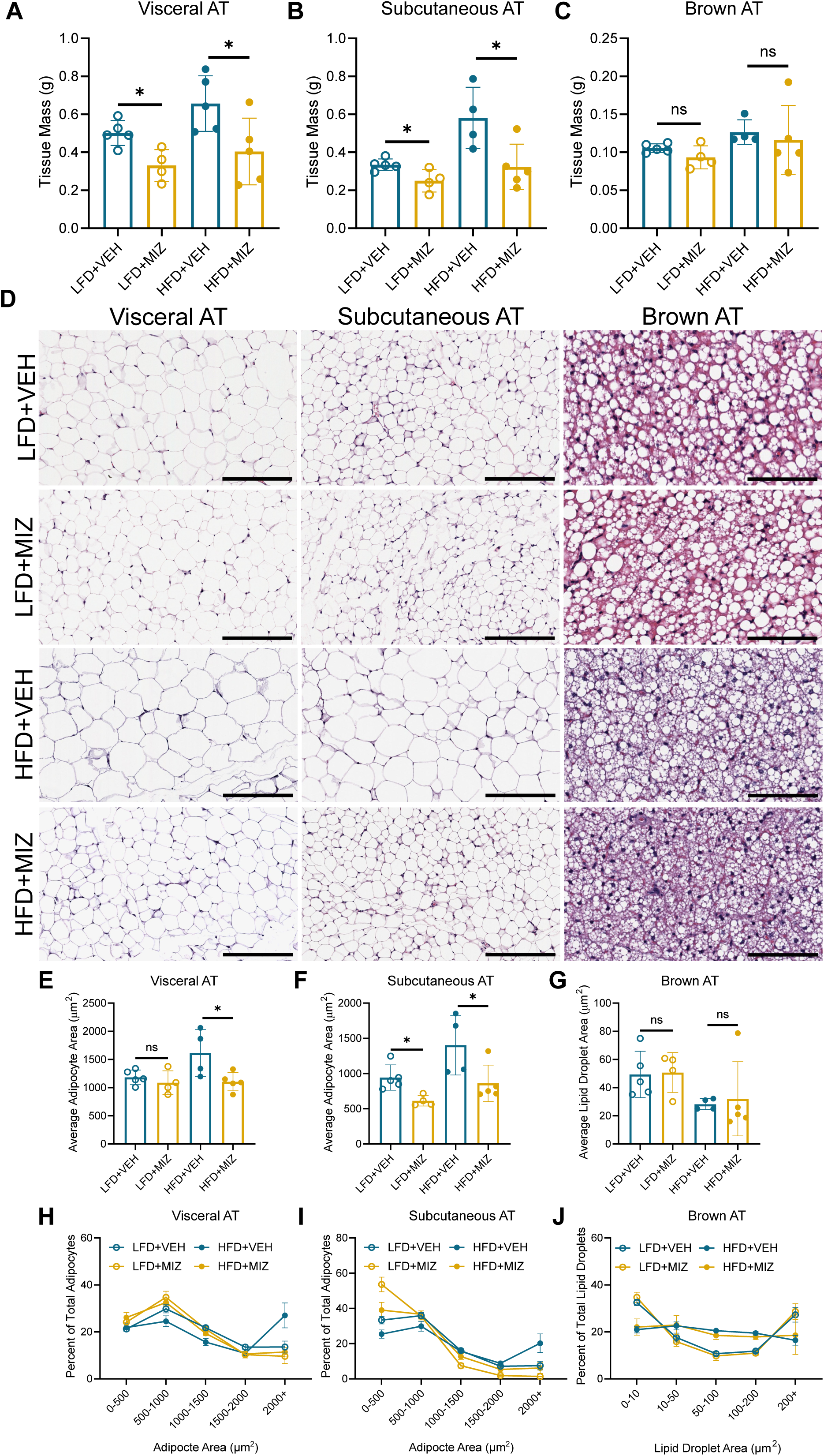
Mizoribine treatment promotes reduction in adipose tissue mass and adipocyte size. (A – C) Terminal tissue mass of visceral adipose tissue (A, VAT), subcutaneous adipose tissue (B, SAT), and brown adipose tissue (C, BAT) from male mice treated for 30 days with a low fat diet (LFD) or high fat diet (HFD) and daily injections of mizoribine (MIZ) or vehicle (VEH). N = 4-5 per group. Error bars represent mean ± standard deviation. Significance indicative of two-tailed student’s *t*-test. Data representative of at least 3 independent experimental cohorts. (D) Representative images of hematoxylin and eosin (H&E) stained sections of VAT, SAT, and BAT from male mice treated with a LFD or HFD and MIZ or VEH injections for 30 days. Scale bars = 200 µm for VAT and SAT and scale bars = 100 µm for BAT. (E-G) Quantification of visceral adipocyte (E), subcutaneous adipocyte (F), and brown adipose lipid droplet (G) size in male mice treated for 30 days with a LFD or HFD and MIZ or VEH. N = 4-5 per group. Error bars represent mean ± standard deviation. Significance indicative of two-tailed student’s *t*-test. (H-J) Frequency distribution of visceral adipocyte (H), subcutaneous adipocyte (I), and brown adipose lipid droplet (J) size in male mice treated for 30 days with a LFD or HFD and MIZ or VEH. N = 4-5 per group. Data representative of at least 2 independent experimental cohorts. ns > 0.05, ∗ *p* ≤ 0.05, ∗∗ *p* ≤ 0.01, ∗∗∗ *p* ≤ 0.001, and ∗∗∗∗ *p* ≤ 0.0001.

### 3.3 Mizoribine does not affect lipid absorption

To evaluate the effects of mizoribine treatment on nutrient absorption, we examined the fecal free fatty acid (FFA) profile of mice treated with mizoribine or vehicle injections and placed on a HDF (Fig. 3 A). Neither the concentration nor the composition of FFAs in the feces was significantly different between the two treatment groups, indicating that nutrient absorption was unaffected by the inhibition of purine biosynthesis (Fig. 3 B – D).

**Figure 3.**
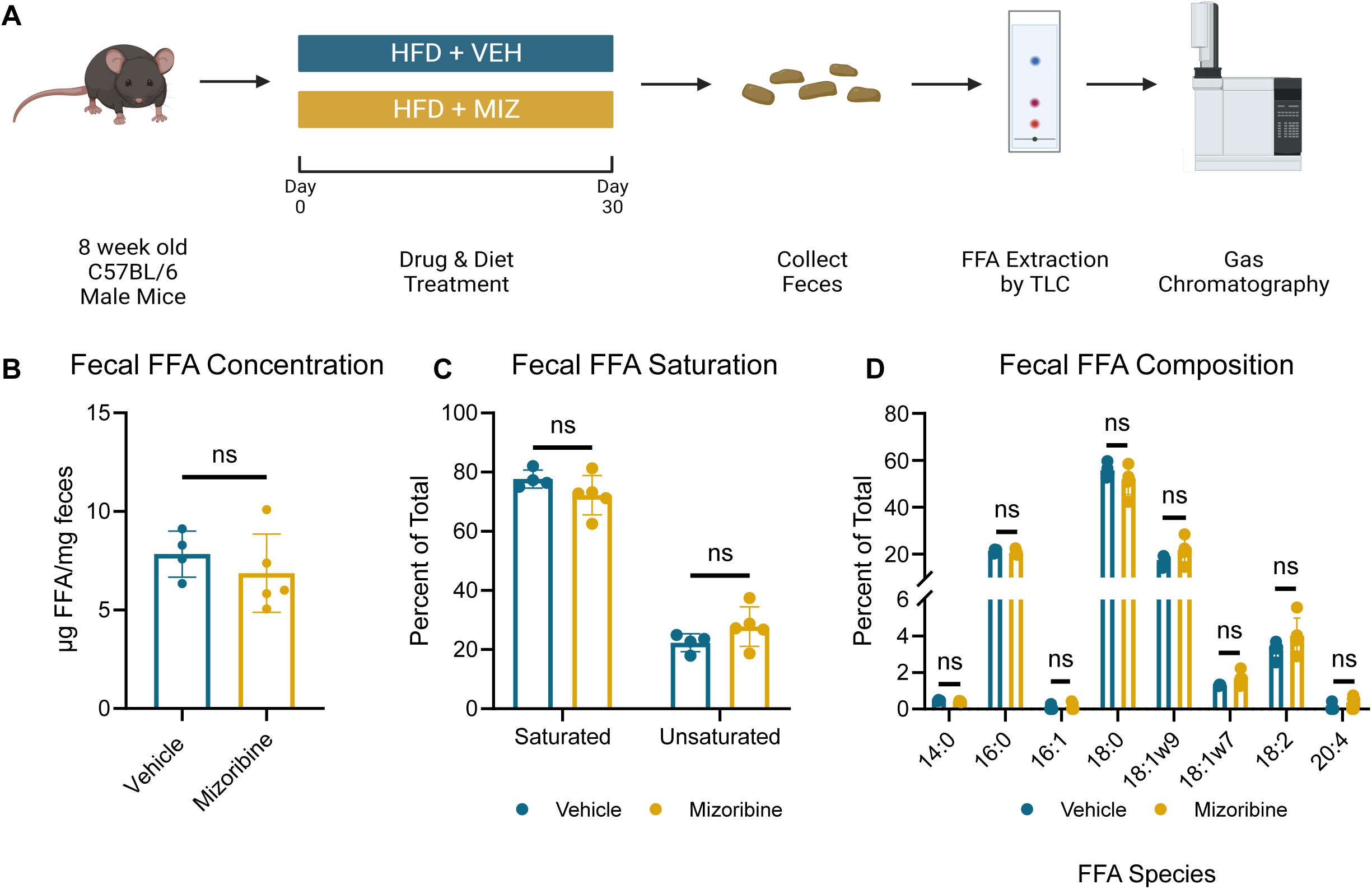
Mizoribine-treated mice do not experience changes in nutrient absorption. (A) Schematic of the experimental setup. Free fatty acid concentration (B), percent saturation (C), and relative species abundance (D) from the feces of male mice after being treated with a high fat diet and vehicle or mizoribine injections for 30 days. N = 5. Error bars represent mean ± standard deviation. For concentration (B) and percent saturation (C), significance indicative of two-tailed student’s *t*-test. For composition (D), significance indicative of Holm-Sidak corrected two-tailed *t*-test. ns > 0.05, ∗ *p* ≤ 0.05, ∗∗ *p* ≤ 0.01, ∗∗∗ *p* ≤ 0.001, and ∗∗∗∗ *p* ≤ 0.0001.

### 3.4 Mizoribine-treated mice eat less

Given that many anti-obesity medications work by suppressing appetite and stimulating satiation, we investigated changes in food consumption and animal behavior [9]. After 30 days, mizoribine-treated mice had eaten less in total (Fig. 4 A); however, circulating leptin levels were not increased in the mizoribine group, suggesting leptin-independent mechanisms of appetite suppression (Fig. 4 B). Water consumption was not significantly altered (Fig. 4 C). To assess the possibility of mizoribine-induced malaise, we measured conditioned taste aversion between the vehicle and mizoribine groups (Fig. 4 D). Mice treated with mizoribine did not develop any significant, learned association specific to mizoribine (Fig. 4 E, S3 A – D). Taken together, these results suggest that while mizoribine suppresses food intake, the mechanism is likely independent of drug-induced nausea, as no taste aversion was observed.

**Figure 4.**
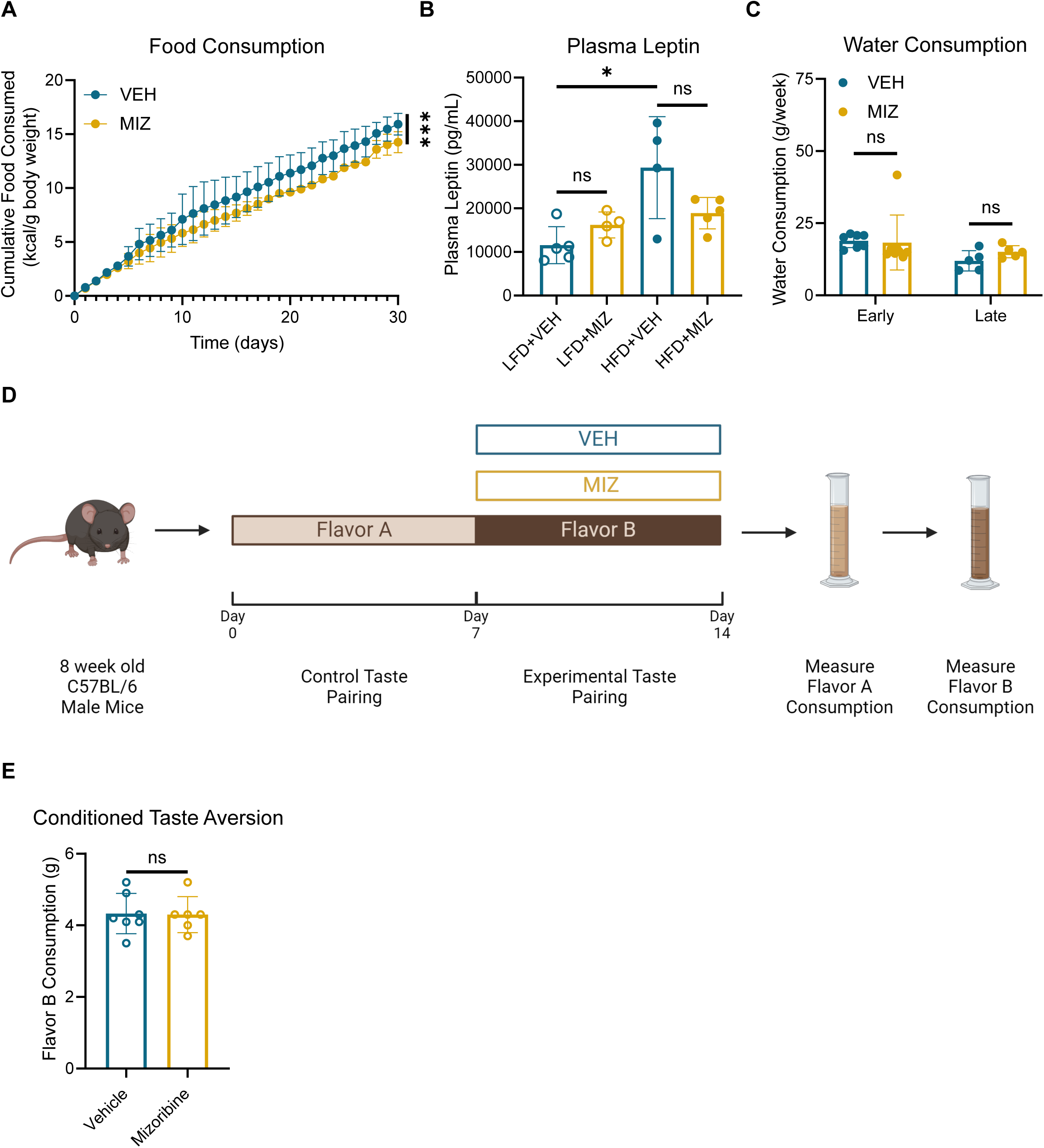
Food consumption is reduced under mizoribine treatment. (A) Cumulative food consumption (normalized to body weight) in male mice treated with a HFD and mizoribine or vehicle. N = 2-3 cages. Error bars represent mean ± standard deviation. Significance indicative of linear regression comparison of slopes. (B) Plasma leptin concentration in male mice after 30 days of treatment with a high or low fat diet and vehicle or mizoribine injections. N = 4-5. Error bars represent mean ± standard deviation. Significance indicative of two-tailed student’s *t*-test. (C) Average weekly water consumption in male mice after being treated with a high fat diet and vehicle or mizoribine injection. Early time point represents consumption during days 0-6 of treatment, late time point represents consumption after 30 days of treatment, during days 30 – 37 of treatment. N = 5-8. Error bars represent mean ± standard deviation. Significance indicative of two-tailed student’s *t*-test. (D) Schematic of the conditioned taste aversion experimental setup. (E) Measurement of flavor B consumption on day 16 of the experiment, comparing conditioned taste aversion to flavor B associated with mizoribine versus vehicle treatment. N = 6-7. Error bars represent mean ± standard deviation. Significance indicative of two-tailed student’s *t*-test. ns > 0.05, ∗ *p* ≤ 0.05, ∗∗ *p* ≤ 0.01, ∗∗∗ *p* ≤ 0.001, and ∗∗∗∗ *p* ≤ 0.0001.

### 3.5 Mizoribine-treated mice do not expend more energy

Although most clinically used anti-obesity drugs aim to reduce energy uptake, enhancement of energy expenditure has also been tested as a strategy for counteracting caloric surplus [10,40]. We examined whether mizoribine had an effect on organismal energy expenditure using indirect calorimetry at two time points: early in the experiment (during the first week of drug and diet treatment) and late in the experiment (after 30 days of drug and diet treatment). Respiratory dynamics and energy expenditure were not changed early in the experiment (Fig. 5 A – D). However, at these early time points, locomotion was reduced in mizoribine-treated mice, potentially suggesting that other factors may be involved in the maintenance of energy expenditure (Fig. 5 E). At later time points, O2 consumption, CO2 production, and the total RER were not different between control and mizoribine-treated mice (Fig. 5 F – H). However, during the dark cycle, when the mice are active, mizoribine increased RER (Fig. 5 H). This increase in RER is likely due to the reduced fat mass of mizoribine-treated mice which promotes their reliance on carbohydrates for energy (Fig. 5 H). At the late time point, energy expenditure was reduced in mizoribine-treated mice, consistent with lower body mass of this group; however, locomotion was unaltered at this later time point (Fig. 5 I – J). The discrepancy between the reduced locomotion and maintained energy expenditure in mizoribine-treated mice at early time points may indicate activation of other mechanisms such as an increased basal metabolic rate or increased thermogenesis.

**Figure 5.**
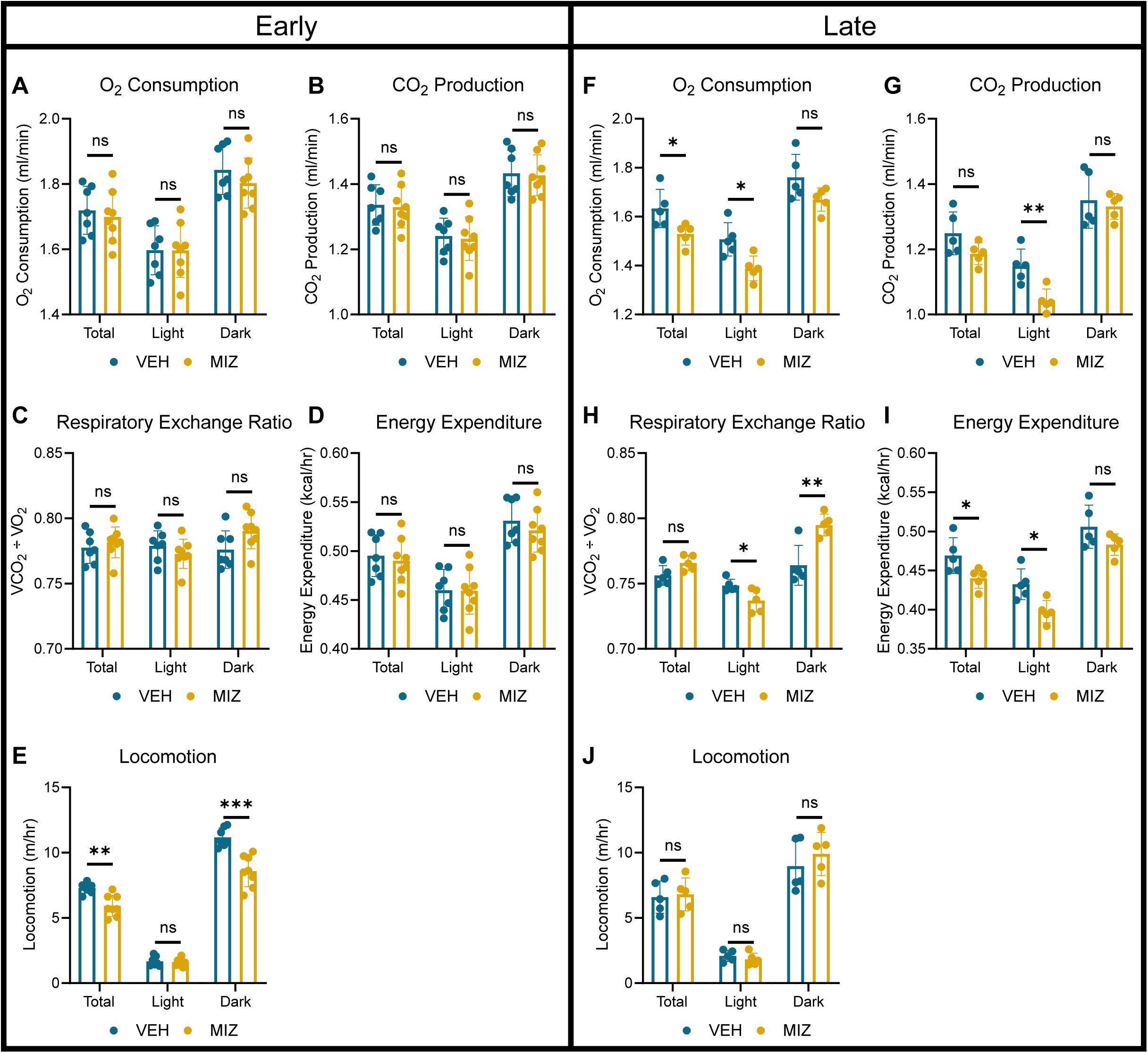
Indirect Calorimetry reveals that energy expenditure is not elevated under mizoribine treatment. (A – E) Oxygen consumption rates (A), carbon dioxide production rates (B), respiratory exchange ratios (C), energy expenditure (D), and locomotion (E) in mizoribine- and vehicle-treated mice fed a high fat diet during the first week of treatment during the full day and light and dark photoperiods. (F – J) Oxygen consumption rates (F), carbon dioxide production rates (G), respiratory exchange ratios (H), energy expenditure (I), and locomotion (J) in mizoribine- and vehicle-treated mice fed a high fat diet after 30 days of treatment during the full day and light and dark photoperiods. N = 5-8. Error bars represent mean ± standard deviation. Significance indicative of two-tailed student’s *t*-test. ns *p* > 0.05, ∗ *p* ≤ 0.05, ∗∗ *p* ≤ 0.01, ∗∗∗ *p* ≤ 0.001.

### 3.6 Thermogenesis is increased in mizoribine-treated mice

Because the mizoribine-treated mice maintained energy expenditure while exhibiting reduced activity, we examined thermogenic processes in these mice. We found that over 30 days of drug and diet treatment, body temperature was elevated in mizoribine-treated mice (Fig. 6 A – B, S4 E). Likewise, we found that mizoribine-treated mice were more tolerant to acute cold exposure (Fig. 6 C). This led us to investigate several thermogenic futile cycles as potential mechanisms underlying mizoribine-induced protection against weight gain. Although uncoupling protein 1 (UCP1)-mediated thermogenesis in BAT is thought to be the primary pathway for non-shivering heat generation in mammals, we did not observe changes in *Ucp1* mRNA or protein expression in BAT (Fig. 6 D – E, S4 D). Furthermore, PR domain containing 16 (*Prdm16)*, a transcriptional regulator of brown fat determination, and peroxisome proliferator-activated receptor γ coactivator 1α (*Ppargc1a)*, a critical regulator of mitochondrial biogenesis and BAT thermogenesis [41,42], were unaltered by the mizoribine treatment (Fig. S4 D). In subcutaneous white adipose tissue, *Ucp1* and *Ppargc1a* expression were not increased by mizoribine, suggesting that there was no beiging of the tissue (Fig. S4 A, C). Markers of lipolysis and fatty acid oxidation were not altered in the adipose tissue of mizoribine-treated mice (Fig. S4 A – B). Finally, expression of creatine kinase B (CKB), another thermogenic futile cycle protein [43], was not changed (Fig. 6 D – E, S4 A). Muscle tissue futile cycles have also been characterized as an alternative to UCP1-mediated thermogenesis [44]. For example, sarcolipin (SLN) is a transmembrane polypeptide that reduces the efficiency of the sarcoplasmic reticulum Ca^2+^-ATPase (SERCA). When SLN is bound to SERCA, ATP hydrolysis is uncoupled from SERCA-mediated Ca^2+^ transport into the SR, resulting in futile cycling and heat production in muscle [45]. We found that expression of SLN was increased in the soleus and diaphragm of mizoribine-treated mice (Fig. 6 F, I, S5 G – H) Previous studies have shown that thyroid hormones downregulate SLN expression [46,47] and that hypothyroid mice exhibit elevated thermogenesis owing to induction of SLN [32,48], which prompted us to quantitate thyroid hormone levels. Surprisingly, in mizoribine-treated mice, plasma concentrations of the thyroid hormone tri-iodothyronine (T_3_) remained unchanged, whereas thyroxine (T_4_) concentrations were elevated (Fig. 6 G – H). These results indicate that mizoribine may repress sarcolipin expression by thyroid hormone-independent mechanisms or by tissue-specific downregulation of thyroid hormone action, independent of plasma thyroid hormone levels.

**Figure 6.**
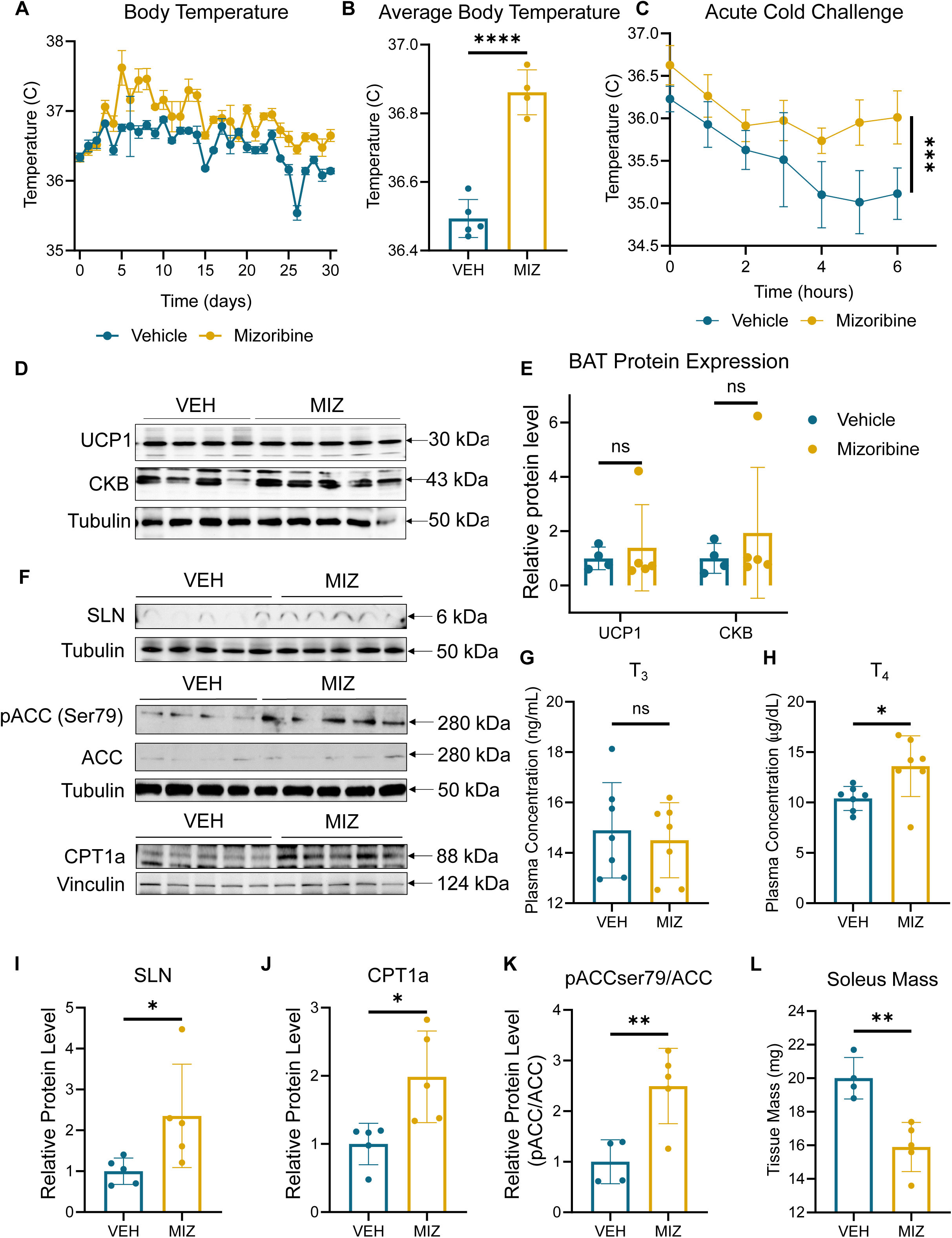
Mizoribine treatment is associated with UCP1 independent thermogenesis. (A) Daily body temperature (as detected by infrared thermometry) in male mice treated with a high fat diet and mizoribine or vehicle for 30 days. N = 4-5. Error bars represent mean ± standard error of the mean. (B) Average body temperature in male mice treated with a high fat diet and mizoribine or vehicle for 30 days. N = 4-5. Error bars represent mean ± standard deviation. Significance indicative of two-tailed student’s *t*-test. (C) Acute cold exposure challenge in male mice treated with a high fat diet and mizoribine or vehicle for 30 days. N = 7-8. Error bars represent mean ± standard deviation. Significance indicative of linear regression comparison of slopes. (D) Immunoblot analysis of UCP1, CKB, and Tubulin in brown adipose tissue from male mice fed a high fat diet and treated with injections of vehicle or mizoribine. (E) Quantification of UCP1 and CKB (Normalized to Tubulin) in brown adipose tissue from male mice fed a high fat diet and treated with injections of vehicle or mizoribine. N = 4-5. Error bars represent mean ± standard deviation. Significance indicative of two-tailed student’s *t*-test. (F) Immunoblot analysis of SLN, p-ACC (Ser79), ACC, CPT1a, Tubulin, and Vinculin in soleus muscle tissue from male mice fed a high fat diet and treated with injections of vehicle or mizoribine. (G – H) Plasma concentration of thyroid hormones T_3_ (G) and T_4_ (H) from male mice treated with a high fat diet and mizoribine or vehicle for 30 days. N = 7. Error bars represent mean ± standard deviation. Significance indicative of two-tailed student’s *t*-test. (I – K) Quantification of soleus immunoblots in F. SLN normalized to Tubulin (I), CPT1a normalized to Vinculin (J), and pACC/ACC ratio (K). N = 4-5. Error bars represent mean ± standard deviation. Significance indicative of two-tailed student’s *t*-test. (L) Soleus muscle mass from male mice fed a high fat diet and treated with injections of vehicle or mizoribine. N = 4-5. Error bars represent mean ± standard deviation. Significance indicative of two-tailed student’s *t*-test. ns > 0.05, ∗ *p* ≤ 0.05, ∗∗ *p* ≤ 0.01, ∗∗∗ *p* ≤ 0.001, and ∗∗∗∗ *p* ≤ 0.0001.

Overexpression of SLN increases energy expenditure and fat oxidation [49]. In agreement with these studies, we found that expression of fatty acid oxidation enzyme carnitine palmitoyl transferase I (CPT1a) was increased, as was the ratio of phosphorylated to unphosphorylated acetyl-CoA carboxylase (pACC:ACC) by the mizorbine treatment, indicative of increased fatty acid oxidation (Fig. 6 F, J – K). No changes in the expression of SLN, pACC, and ACC were observed in other muscles (Fig. S5 D – F). Correspondingly, muscle tissue mass was reduced in the soleus but unaltered in other muscles (Fig. 6 L, S5 A – C). These studies indicate that inhibition of purine biosynthesis may stimulate thermogenesis and energy expenditure in the soleus muscle by activating the sarcolipin-mediated SERCA futile cycle.

### 3.7 Mizoribine treatment protects against obesity-mediated metabolic dysfunction

Diet-induced obesity is commonly associated with impaired insulin sensitivity and glucose tolerance and elevated fasting blood glucose levels. Mizoribine treatment reduced basal fasting blood glucose levels (Fig. 7 A). To determine the effects of mizoribine on glucose metabolism, we performed intraperitoneal glucose tolerance tests. Blood glucose clearance was accelerated in mizoribine-treated mice (Fig. 7 B – C). In addition, the plasma insulin levels were lower after the glucose injection (Fig. 7 D). Immunofluorescence analysis of pancreatic islet cells showed that mizoribine treatment did not impact pancreatic islet endocrine cell area, suggesting that mizoribine does not affect the pancreatic islets on a structural level (Fig. 7 E, S6 A). Taken together, these results suggest that mizoribine treatment improves glucose tolerance in mice fed a HFD.

**Figure 7.**
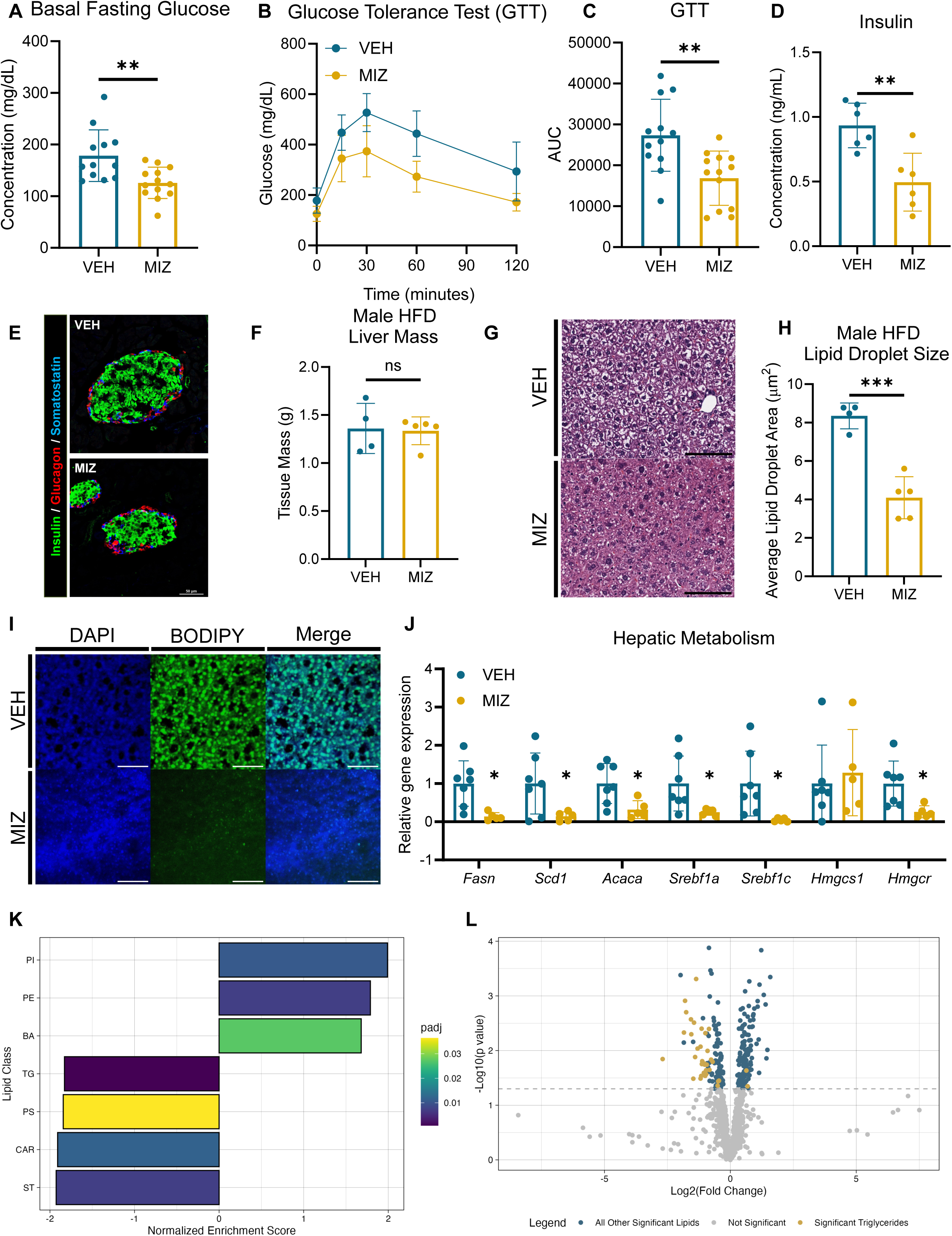
Mizoribine treatment improves systemic metabolic health. (A) Basal fasting glucose in male mice treated with a high fat diet and mizoribine or vehicle injections for 30 days. N = 12-13. Error bars represent mean ± standard deviation. Significance indicative of two-tailed student’s *t*-test. (B – C) Intraperitoneal glucose tolerance test on male mice treated with a high fat diet and mizoribine or vehicle injections for 30 days. Glucose concentration over time (B) and baseline-adjusted area under the curve quantification (C) of the glucose versus time curves. N = 12-13. Error bars represent mean ± standard deviation. Significance indicative of two-tailed student’s *t*-test. (D) Plasma insulin concentration during glucose tolerance test (15 minutes after the dextrose injection). N = 6. Error bars represent mean ± standard deviation. Significance indicative of two-tailed student’s *t*-test. (E) Representative images of insulin (green), glucagon (red), and somatostatin (blue) labeled pancreatic islets. (F) Terminal liver mass from male mice treated for 30 days with a high fat diet (HFD) and daily injections of mizoribine (MIZ) or vehicle (VEH). N = 4-5 per group. Error bars represent mean ± standard deviation. Significance indicative of two-tailed student’s *t*-test. (G) Representative images of hematoxylin and eosin (H&E) stained sections of liver from male mice treated with a HFD and either vehicle or mizoribine injections. Scale bars = 100 µm (H) Quantification of hepatic lipid droplet size in male mice with a HFD and either vehicle or mizoribine injections. N = 4-5 per group. Error bars represent mean ± standard deviation. Significance indicative of two-tailed student’s *t*-test. (I) Representative images of DAPI and BODIPY stained sections of liver from male mice treated with a HFD and either vehicle or mizoribine injections. Scale bars = 50 µm. (J) Gene expression analysis of *Fasn, Scd1, Acaca, Srebf1a, Srebf1c, Hmgcr,* and *Hmgcs1* (normalized to *Rps16*). N = 5-7 per group. Error bars represent mean ± standard deviation. Significance indicative of two-tailed student’s *t*-test. ∗ *p* ≤ 0.05. (K) Lipid sets with significant (padj < 0.05) differential enrichment in the plasma of vehicle- and mizoribine-treated male mice. “padj” represents the FDR-adjusted p-value. Negative normalized enrichment scores represent a decrease in mizoribine samples compared to vehicle, while positive values represent an increase in mizoribine samples. Lipid classes: PI – phosphatidylinositol, PE – phosphatidylethanolamine, BA – bile acid, TG – triglyceride, PS – phosphatidylserine, CAR – acylcarnitine, ST – stigmasterol. (L) Volcano plot of differentially abundant lipids in the plasma of vehicle- and mizoribine-treated male mice. Significant (p < 0.05), differentially abundant triglyceride species are shown in gold. All other significantly differentially expressed lipid species are shown in teal, and nonsignificant lipid species are shown in gray. ns > 0.05, ∗ *p* ≤ 0.05, ∗∗ *p* ≤ 0.01, ∗∗∗ *p* ≤ 0.001, and ∗∗∗∗ *p* ≤ 0.0001.

Considering the comorbidity of obesity and metabolic-associated fatty liver disease, we investigated changes in liver metabolism and hepatic lipid accumulation. While liver mass was constant (Fig. 7 F), ectopic lipid accumulation in the liver was reduced under mizoribine treatment in HFD-fed male mice (Fig. 7 G – I). Lipid accumulation in the livers of LFD-fed male mice was unchanged under mizoribine treatment (Fig. S6 B – D). Likewise, expression of genes associated with lipid storage, *Fasn, Scd1, Acaca, Srebf1a,* and *Srebp1c* was reduced in mizoribine-treated mice (Fig. 7 J). Expression of *Hmgcr*, a key gene involved in cholesterol biosynthesis, was also reduced (Fig. 7 I). To further investigate changes in lipid metabolism, we analyzed the circulating plasma lipidome. Mizoribine-treatment significantly altered the lipidome (Fig. 7 K, S6 E). In particular, circulating triglyceride levels decreased in mizoribine-treated mice (Fig. 7 K – L, S6 F). This was largely driven by lower levels of circulating triglycerides with greater degrees of unsaturation (Fig. S6 F). These data suggest that mizoribine-treated mice, despite their reduced adipose storage, are not lipodystrophic and do not show signs of elevated ectopic lipid accumulation. Rather, mizoribine-treated mice have improved glucose tolerance and appear to be metabolically healthier than control mice when challenged with a HFD.

## 4. Discussion

In this study, we have demonstrated that pharmacological inhibition of IMPDH1/2, key enzymes in the *de novo* purine biosynthesis pathway, effectively mitigates HFD-induced weight gain in mice. Notably, mizoribine did not significantly affect weight when mice were fed a standard LFD, underscoring its potential specificity towards HFD conditions. Mizoribine treatment prevented fat mass from accumulating, without significantly altering lean mass. This aligns with previous observations made using another IMPDH inhibitor, mycophenolate mofetil, suggesting a class effect among purine biosynthesis inhibitors in the context of obesity [50].

We next considered the mechanism by which purine biosynthesis inhibition prevents HFD-mediated weight gain. Body weight gain is a consequence of positive energy balance. Energy intake, nutrient absorption, and energy expenditure are key determinants of energy balance. The mechanism by which mizoribine regulates adiposity appears to be multifaceted. Mizoribine treatment did not alter lipid content in the feces, establishing that nutrient absorption is not part of the mechanism by which purine inhibition protects mice against weight gain. Mizoribine decreased food intake; however, our studies indicated that a leptin-independent pathway may be involved in this regulation, as leptin levels were unchanged. Although mizoribine treatment did not change total energy expenditure, it did reduce the contribution from locomotion whereas thermogenesis was increased, resulting in a net zero effect on energy expenditure. It is unclear whether the increase in thermogenesis results from or causes the decrease in locomotor activity.

We further investigated the mechanism by which mizoribine increases thermogenesis in mice. Previous studies have reported conflicting effects of purine biosynthesis inhibition on UCP1, with one study showing that the IMPDH inhibitor mycophenolic acid suppressed UCP1 activity [51] and another suggesting that mycophenolate mofetil (MMF), a prodrug of MPA, stimulates UCP1 expression and adipose tissue browning [52]. Under our treatment conditions, UCP1 expression did not change, nor did we observe adipose tissue beiging. Creatine kinase B (CKB), another regulator of adipose thermogenesis [53,54], was also not significantly upregulated by the mizoribine treatment in BAT. However, the expression of SLN, a regulator of muscle thermogenesis, was increased in skeletal muscle after mizoribine treatment. Previously published work demonstrated that loss of SLN promotes weight gain [44,49] establishing the role of non-shivering thermogenesis in energy balance. Thus, it is likely that the increase in SLN we observed is at least a part of the mechanism by which mizoribine impacts adiposity and organismal physiology. Moreover, it has been shown that, in mice that overexpress SLN, the circulating lipid profile is also improved, as demonstrated by their lower triglyceride levels [43]. We observed a similar reduction in levels of plasma triglycerides, particularly those with high-carbon content, after mizoribine treatment, consistent with a preference for nutrient utilization over storage. Interestingly, unsaturated triglyceride levels were preferentially reduced by the mizoribine treatment, which may indicate selective mobilization of these lipids and their utilization for energy. Finally, how SLN is regulated by mizoribine remains unclear. Although we ruled out a decrease in circulating thyroid hormone as a factor in sarcolipin upregulation in the muscle, the possibility remains that the increase in thermogenesis was driven by tissue-specific thyroid hormone down-regulation [48].

As adipocytes reach their maximal capacity or become dysfunctional, triglycerides and other lipid species may accumulate in the liver and skeletal muscle, resulting in insulin resistance [55]. The inability to store lipid in adipose tissue manifests clinically as a heterogeneous class of disorders known as lipodystrophies [56]. Lipodystrophies often present with severe insulin resistance from ectopic lipid storage—and, indeed, the adipose storage inefficiencies observed in lipodystrophies have been posited to arise from mechanisms partially parallel to those that give rise to them in obesity [57]. Thus, we investigated the effects of purine inhibition on metabolic health. We found that mizoribine improves systemic glucose metabolism and reduces ectopic lipid accumulation strongly arguing against lipodystrophy.

Nucleotide biosynthesis inhibitors have been used clinically to treat a variety of malignancies [58]. Low doses of nucleotide biosynthesis inhibitors are also used as immunosuppressant agents to treat a number of autoimmune conditions, including systemic lupus erythematosus, rheumatoid arthritis, ulcerative colitis, and Crohn’s disease [59–62]. Similarly, nucleotide biosynthesis inhibitors are widely used to prevent rejection of solid organ transplants [63–65]. Given that nucleotide biosynthesis inhibitors are widely used in various clinical settings, our findings also raise important considerations pertaining to their broader metabolic impacts. Thus, purine inhibitors may improve systemic metabolism in obese or overweight patients.

Taken together, our results reveal an important physiological connection between purine availability and adiposity. Future investigations should aim to delineate the precise molecular mechanisms by which purine biosynthesis influences thermogenesis and food intake. It will also be worthwhile to further test the hypothesis that mizoribine hyperactivates slow-twitch oxidative muscle tissues (soleus and diaphragm) in which SLN is mainly expressed, to consume fatty acids, thus diverting lipids away from the adipose tissue and the liver and preventing them from being stored there. Insights into these mechanisms could bring to light new therapeutic targets that could be used in combating obesity and related metabolic disorders.

## Data Availability

Any information required to reanalyze the data reported in this article is available from the lead contact upon request.

## Acknowledgments

E.Z. was supported by a P30 058404 DDRTC Pilot and Feasibility Award, the DK020593 Vanderbilt Diabetes and Research Training Center, the Vanderbilt University Seeding Success grant, a AHA Career Development Award 23CDA1050768 and NIGMS R35GM154684. S.H.P was supported by a 1IK2BX005401 grant from the Department of Veterans Affairs and funds from the Department of Medicine at Vanderbilt University Medical Center. T.T.H was supported by funds from the Department of Medicine at Vanderbilt University Medical Center. M.R.M. was supported by the Howard Hughes Medical Institute Hanna H. Gray Fellows Program Faculty Phase (Grant# GT15655) and the Burroughs Welcome Fund PDEP Transition to Faculty (Grant# 1022604). Histological sectioning and staining were performed by the Vanderbilt Translational Pathology Shared Resource and was supported by NIH/NCI Cancer Center Support Grant 5P30 CA68485-19. Histological imaging was performed using the Vanderbilt Islet and Pancreas Analysis Core and was supported by the Vanderbilt Diabetes Research and Training Center (NIH grant DK20593). Confocal imaging was performed using the Vanderbilt Cell Imaging Shared Resource (CISR) and was supported by NIH grants 1S10OD012324-01, 1S10OD021630-01, CA68485, DK20593, DK58404, DK59637 and EY08126. Energy expenditure experiments were performed with assistance from the Vanderbilt Mouse Metabolic Phenotyping Center and were supported by NIH grant 5U2C DK059637. Gas chromatography and radioimmunoassay experiments were performed by the Vanderbilt Analytical Services Core and were supported by NIH grant DK020593. We would also like to acknowledge the Huck Institutes’ Metabolomics Core Facility (RRID:SCR_023864) for use of the OE 240 LCMS and Sergei Koshkin for helpful discussions on sample preparation and analysis. We thank Jose Maldonado, Kathleen DelGiorno, Katherine Ankenbauer, and the Vanderbilt Neurovisualization core for the use of the Axios Scan.Z1. Figures 1 A – B, 3 A, and 4 D were created using BioRender.com. The authors thank Mila Lazarević, Genesis Wilson, Megan Altemus, Merrygay James, and Carlo Malabanan for assistance and advice.

## Author contributions

E.Z. conceived the study. E.Z. and J.W.M. wrote the manuscript, and all authors contributed to editing. J.W.M., W.Y.P., A.M.E., A.B.S., and M.Z.L. designed experiments. J.W.M., W.Y.P., and A.M.E. performed formal analysis. J.W.M., W.Y.P., A.M.E., J.A.P., C.M., Z.S., J.J.R., and G.N.R performed experiments on gene expression, metabolite, hormone, and protein abundance, and microscopy analysis. P.P., A.C.M., M.R.M., and J.W.M. performed lipidomic analysis. M.Z.L., T.T.H., N.C.W., S.H.P., K.C.C., R.W.S., N.C., L.L., E.S.C., M.R.M., and E.Z. provided scientific guidance, resources, and supervision.

## Conflict of interest

The authors declare that they have no conflict of interest.

## Notes

https://github.com/jacobwm/Mizoribine-Lipidomics

### Competing Interest Statement

The authors have declared no competing interest.

## References

[1] Bryan, S., Afful, J., Carroll, M., Te-Ching, C., Orlando, D., Fink, S., et al., 2021. NHSR 158. National Health and Nutrition Examination Survey 2017–March 2020 Pre-pandemic Data Files. National Center for Health Statistics (U.S.).

[2] Ward, Z.J., Bleich, S.N., Cradock, A.L., Barrett, J.L., Giles, C.M., Flax, C., et al., 2019. Projected U.S. State-Level Prevalence of Adult Obesity and Severe Obesity. New England Journal of Medicine 381(25): 2440–50, Doi: 10.1056/NEJMsa1909301.

[3] The GBD 2015 Obesity Collaborators, 2017. Health Effects of Overweight and Obesity in 195 Countries over 25 Years. New England Journal of Medicine 377(1): 13–27, Doi: 10.1056/NEJMoa1614362.

[4] Haffner, S.M., 1990. Cardiovascular Risk Factors in Confirmed Prediabetic Individuals: Does the Clock for Coronary Heart Disease Start Ticking Before the Onset of Clinical Diabetes? JAMA 263(21): 2893, Doi: 10.1001/jama.1990.03440210043030.

[5] Blüher, M., 2019. Obesity: global epidemiology and pathogenesis. Nature Reviews Endocrinology 15(5): 288–98, Doi: 10.1038/s41574-019-0176-8.

[6] Powell-Wiley, T.M., Poirier, P., Burke, L.E., Després, J.-P., Gordon-Larsen, P., Lavie, C.J., et al., 2021. Obesity and Cardiovascular Disease: A Scientific Statement From the American Heart Association. Circulation 143(21), Doi: 10.1161/CIR.0000000000000973.

[7] ElSayed, N.A., Aleppo, G., Aroda, V.R., Bannuru, R.R., Brown, F.M., Bruemmer, D., et al., 2023. 8. Obesity and Weight Management for the Prevention and Treatment of Type 2 Diabetes: *Standards of Care in Diabetes—*2023. Diabetes Care 46(Supplement_1): S128–39, Doi: 10.2337/dc23-S008.

[8] Tak, Y.J., Lee, S.Y., 2021. Long-Term Efficacy and Safety of Anti-Obesity Treatment: Where Do We Stand? Current Obesity Reports 10(1): 14–30, Doi: 10.1007/s13679-020-00422-w.

[9] Idrees, Z., Cancarevic, I., Huang, L., 2022. FDA-Approved Pharmacotherapy for Weight Loss Over the Last Decade. Cureus, Doi: 10.7759/cureus.29262.

[10] Abdi Beshir, S., Ahmed Elnour, A., Soorya, A., Parveen Mohamed, A., Sir Loon Goh, S., Hussain, N., et al., 2023. A narrative review of approved and emerging anti-obesity medications. Saudi Pharmaceutical Journal: SPJ: The Official Publication of the Saudi Pharmaceutical Society 31(10): 101757, Doi: 10.1016/j.jsps.2023.101757.

[11] Hepler, C., Gupta, R.K., 2017. The expanding problem of adipose depot remodeling and postnatal adipocyte progenitor recruitment. Molecular and Cellular Endocrinology 445: 95–108, Doi: 10.1016/j.mce.2016.10.011.

[12] Sun, K., Kusminski, C.M., Scherer, P.E., 2011. Adipose tissue remodeling and obesity. The Journal of Clinical Investigation 121(6): 2094–101, Doi: 10.1172/JCI45887.

[13] Sun, K., Tordjman, J., Clément, K., Scherer, P.E., 2013. Fibrosis and adipose tissue dysfunction. Cell Metabolism 18(4): 470–7, Doi: 10.1016/j.cmet.2013.06.016.

[14] Klöting, N., Blüher, M., 2014. Adipocyte dysfunction, inflammation and metabolic syndrome. Reviews in Endocrine & Metabolic Disorders 15(4): 277–87, Doi: 10.1007/s11154-014-9301-0.

[15] Gonzalez, F.J., Xie, C., Jiang, C., 2019. The role of hypoxia-inducible factors in metabolic diseases. Nature Reviews Endocrinology 15(1): 21–32, Doi: 10.1038/s41574-018-0096-z.

[16] Fontana, L., Cummings, N.E., Arriola Apelo, S.I., Neuman, J.C., Kasza, I., Schmidt, B.A., et al., 2016. Decreased Consumption of Branched-Chain Amino Acids Improves Metabolic Health. Cell Reports 16(2): 520–30, Doi: 10.1016/j.celrep.2016.05.092.

[17] Wang, T.J., Larson, M.G., Vasan, R.S., Cheng, S., Rhee, E.P., McCabe, E., et al., 2011. Metabolite profiles and the risk of developing diabetes. Nature Medicine 17(4): 448–53, Doi: 10.1038/nm.2307.

[18] Park, S., Sadanala, K.C., Kim, E.-K., 2015. A Metabolomic Approach to Understanding the Metabolic Link between Obesity and Diabetes. Molecules and Cells 38(7): 587–96, Doi: 10.14348/molcells.2015.0126.

[19] Petrus, P., Lecoutre, S., Dollet, L., Wiel, C., Sulen, A., Gao, H., et al., 2020. Glutamine Links Obesity to Inflammation in Human White Adipose Tissue. Cell Metabolism 31(2): 375–390.e11, Doi: 10.1016/j.cmet.2019.11.019.

[20] Tsushima, Y., Nishizawa, H., Tochino, Y., Nakatsuji, H., Sekimoto, R., Nagao, H., et al., 2013. Uric Acid Secretion from Adipose Tissue and Its Increase in Obesity. Journal of Biological Chemistry 288(38): 27138–49, Doi: 10.1074/jbc.M113.485094.

[21] Cummins, T.D., Holden, C.R., Sansbury, B.E., Gibb, A.A., Shah, J., Zafar, N., et al., 2014. Metabolic remodeling of white adipose tissue in obesity. American Journal of Physiology-Endocrinology and Metabolism 307(3): E262–77, Doi: 10.1152/ajpendo.00271.2013.

[22] Tamba, S., Nishizawa, H., Funahashi, T., Okauchi, Y., Ogawa, T., Noguchi, M., et al., 2008. Relationship between the serum uric acid level, visceral fat accumulation and serum adiponectin concentration in Japanese men. Internal Medicine (Tokyo, Japan) 47(13): 1175–80, Doi: 10.2169/internalmedicine.47.0603.

[23] Johnson, R.J., Nakagawa, T., Sanchez-Lozada, L.G., Shafiu, M., Sundaram, S., Le, M., et al., 2013. Sugar, Uric Acid, and the Etiology of Diabetes and Obesity. Diabetes 62(10): 3307–15, Doi: 10.2337/db12-1814.

[24] Sautin, Y.Y., Nakagawa, T., Zharikov, S., Johnson, R.J., 2007. Adverse effects of the classic antioxidant uric acid in adipocytes: NADPH oxidase-mediated oxidative/nitrosative stress. American Journal of Physiology-Cell Physiology 293(2): C584–96, Doi: 10.1152/ajpcell.00600.2006.

[25] Baldwin, W., McRae, S., Marek, G., Wymer, D., Pannu, V., Baylis, C., et al., 2011. Hyperuricemia as a Mediator of the Proinflammatory Endocrine Imbalance in the Adipose Tissue in a Murine Model of the Metabolic Syndrome. Diabetes 60(4): 1258– 69, Doi: 10.2337/db10-0916.

[26] Nakagawa, T., Hu, H., Zharikov, S., Tuttle, K.R., Short, R.A., Glushakova, O., et al., 2006. A causal role for uric acid in fructose-induced metabolic syndrome. American Journal of Physiology-Renal Physiology 290(3): F625–31, Doi: 10.1152/ajprenal.00140.2005.

[27] Schneider, C.A., Rasband, W.S., Eliceiri, K.W., 2012. NIH Image to ImageJ: 25 years of image analysis. Nature Methods 9(7): 671–5, Doi: 10.1038/nmeth.2089.

[28] Bankhead, P., Loughrey, M.B., Fernández, J.A., Dombrowski, Y., McArt, D.G., Dunne, P.D., et al., 2017. QuPath: Open source software for digital pathology image analysis. Scientific Reports 7(1): 16878, Doi: 10.1038/s41598-017-17204-5.

[29] Folch, J., Lees, M., Sloane Stanley, G.H., 1957. A simple method for the isolation and purification of total lipides from animal tissues. The Journal of Biological Chemistry 226(1): 497–509.

[30] Morrison, W.R., Smith, L.M., 1964. PREPARATION OF FATTY ACID METHYL ESTERS AND DIMETHYLACETALS FROM LIPIDS WITH BORON FLUORIDE--METHANOL. Journal of Lipid Research 5: 600–8.

[31] Weir, J.B.D.B., 1949. New methods for calculating metabolic rate with special reference to protein metabolism. The Journal of Physiology 109(1–2): 1–9, Doi: 10.1113/jphysiol.1949.sp004363.

[32] Kaspari, R.R., Reyna-Neyra, A., Jung, L., Torres-Manzo, A.P., Hirabara, S.M., Carrasco, N., 2020. The paradoxical lean phenotype of hypothyroid mice is marked by increased adaptive thermogenesis in the skeletal muscle. Proceedings of the National Academy of Sciences of the United States of America 117(36): 22544–51, Doi: 10.1073/pnas.2008919117.

[33] Dean, E.D., Li, M., Prasad, N., Wisniewski, S.N., Von Deylen, A., Spaeth, J., et al., 2017. Interrupted Glucagon Signaling Reveals Hepatic α Cell Axis and Role for L-Glutamine in α Cell Proliferation. Cell Metabolism 25(6): 1362–1373.e5, Doi: 10.1016/j.cmet.2017.05.011.

[34] Spears, E., Stanley, J.E., Shou, M., Yin, L., Li, X., Dai, C., et al., 2023. Pancreatic islet α cell function and proliferation requires the arginine transporter SLC7A2, Doi: 10.1101/2023.08.10.552656.

[35] Mohamed, A., Molendijk, J., Hill, M.M., 2020. lipidr: A Software Tool for Data Mining and Analysis of Lipidomics Datasets. Journal of Proteome Research 19(7): 2890–7, Doi: 10.1021/acs.jproteome.0c00082.

[36] Butte, N.F., Liu, Y., Zakeri, I.F., Mohney, R.P., Mehta, N., Voruganti, V.S., et al., 2015. Global metabolomic profiling targeting childhood obesity in the Hispanic population. The American Journal of Clinical Nutrition 102(2): 256–67, Doi: 10.3945/ajcn.115.111872.

[37] Shinde, A.B., Nunn, E.R., Wilson, G.A., Chvasta, M.T., Pinette, J.A., Myers, J.W., et al., 2023. Inhibition of nucleotide biosynthesis disrupts lipid accumulation and adipogenesis. Journal of Biological Chemistry: 104635, Doi: 10.1016/j.jbc.2023.104635.

[38] Pinette, J.A., Myers, J.W., Park, W.Y., Bryant, H.G., Eddie, A.M., Wilson, G.A., et al., 2024. Disruption of nucleotide biosynthesis reprograms mitochondrial metabolism to inhibit adipogenesis. Journal of Lipid Research: 100641, Doi: 10.1016/j.jlr.2024.100641.

[39] Huang, K.-P., Ronveaux, C.C., Knotts, T.A., Rutkowsky, J.R., Ramsey, J.J., Raybould, H.E., 2020. Sex differences in response to short-term high fat diet in mice. Physiology & Behavior 221: 112894, Doi: 10.1016/j.physbeh.2020.112894.

[40] Goldgof, M., Xiao, C., Chanturiya, T., Jou, W., Gavrilova, O., Reitman, M.L., 2014. The Chemical Uncoupler 2,4-Dinitrophenol (DNP) Protects against Diet-induced Obesity and Improves Energy Homeostasis in Mice at Thermoneutrality. Journal of Biological Chemistry 289(28): 19341–50, Doi: 10.1074/jbc.M114.568204.

[41] Puigserver, P., Wu, Z., Park, C.W., Graves, R., Wright, M., Spiegelman, B.M., 1998. A cold-inducible coactivator of nuclear receptors linked to adaptive thermogenesis. Cell 92(6): 829–39, Doi: 10.1016/s0092-8674(00)81410-5.

[42] Seale, P., Kajimura, S., Yang, W., Chin, S., Rohas, L.M., Uldry, M., et al., 2007. Transcriptional control of brown fat determination by PRDM16. Cell Metabolism 6(1): 38–54, Doi: 10.1016/j.cmet.2007.06.001.

[43] Rahbani, J.F., Roesler, A., Hussain, M.F., Samborska, B., Dykstra, C.B., Tsai, L., et al., 2021. Creatine kinase B controls futile creatine cycling in thermogenic fat. Nature 590(7846): 480–5, Doi: 10.1038/s41586-021-03221-y.

[44] Bal, N.C., Maurya, S.K., Sopariwala, D.H., Sahoo, S.K., Gupta, S.C., Shaikh, S.A., et al., 2012. Sarcolipin is a newly identified regulator of muscle-based thermogenesis in mammals. Nature Medicine 18(10): 1575–9, Doi: 10.1038/nm.2897.

[45] Smith, W.S., Broadbridge, R., East, J.M., Lee, A.G., 2002. Sarcolipin uncouples hydrolysis of ATP from accumulation of Ca2+ by the Ca2+-ATPase of skeletal-muscle sarcoplasmic reticulum. The Biochemical Journal 361(Pt 2): 277–86, Doi: 10.1042/0264-6021:3610277.

[46] Johann, K., Cremer, A.L., Fischer, A.W., Heine, M., Pensado, E.R., Resch, J., et al., 2019. Thyroid-Hormone-Induced Browning of White Adipose Tissue Does Not Contribute to Thermogenesis and Glucose Consumption. Cell Reports 27(11): 3385–3400.e3, Doi: 10.1016/j.celrep.2019.05.054.

[47] Minamisawa, S., Uemura, N., Sato, Y., Yokoyama, U., Yamaguchi, T., Inoue, K., et al., 2006. Post-transcriptional downregulation of sarcolipin mRNA by triiodothyronine in the atrial myocardium. FEBS Letters 580(9): 2247–52, Doi: 10.1016/j.febslet.2006.03.032.

[48] Gereben, B., Zavacki, A.M., Ribich, S., Kim, B.W., Huang, S.A., Simonides, W.S., et al., 2008. Cellular and molecular basis of deiodinase-regulated thyroid hormone signaling. Endocrine Reviews 29(7): 898–938, Doi: 10.1210/er.2008-0019.

[49] Maurya, S.K., Bal, N.C., Sopariwala, D.H., Pant, M., Rowland, L.A., Shaikh, S.A., et al., 2015. Sarcolipin Is a Key Determinant of the Basal Metabolic Rate, and Its Overexpression Enhances Energy Expenditure and Resistance against Diet-induced Obesity. The Journal of Biological Chemistry 290(17): 10840–9, Doi: 10.1074/jbc.M115.636878.

[50] Su, H., Gunter, J.H., De Vries, M., Connor, T., Wanyonyi, S., Newell, F.S., et al., 2009. Inhibition of inosine monophosphate dehydrogenase reduces adipogenesis and diet-induced obesity. Biochemical and Biophysical Research Communications 386(2): 351–5, Doi: 10.1016/j.bbrc.2009.06.040.

[51] Fromme, T., Kleigrewe, K., Dunkel, A., Retzler, A., Li, Y., Maurer, S., et al., 2018. Degradation of brown adipocyte purine nucleotides regulates uncoupling protein 1 activity. Molecular Metabolism 8: 77–85, Doi: 10.1016/j.molmet.2017.12.010.

[52] Takahashi, H., Tokura, M., Kawarasaki, S., Nagai, H., Iwase, M., Nishitani, K., et al., 2022. Metabolomics reveals inosine 5’-monophosphate is increased during mice adipocyte browning. The Journal of Biological Chemistry 298(10): 102456, Doi: 10.1016/j.jbc.2022.102456.

[53] Rahbani, J.F., Bunk, J., Lagarde, D., Samborska, B., Roesler, A., Xiao, H., et al., 2024. Parallel control of cold-triggered adipocyte thermogenesis by UCP1 and CKB. Cell Metabolism 36(3): 526–540.e7, Doi: 10.1016/j.cmet.2024.01.001.

[54] Rahbani, J.F., Scholtes, C., Lagarde, D.M., Hussain, M.F., Roesler, A., Dykstra, C.B., et al., 2022. ADRA1A–Gαq signalling potentiates adipocyte thermogenesis through CKB and TNAP. Nature Metabolism 4(11): 1459–73, Doi: 10.1038/s42255-022-00667-w.

[55] Perry, R.J., Samuel, V.T., Petersen, K.F., Shulman, G.I., 2014. The role of hepatic lipids in hepatic insulin resistance and type 2 diabetes. Nature 510(7503): 84–91, Doi: 10.1038/nature13478.

[56] Mann, J.P., Savage, D.B., 2019. What lipodystrophies teach us about the metabolic syndrome. The Journal of Clinical Investigation 129(10): 4009–21, Doi: 10.1172/JCI129190.

[57] Vishvanath, L., Gupta, R.K., 2019. Contribution of adipogenesis to healthy adipose tissue expansion in obesity. The Journal of Clinical Investigation 129(10): 4022–31, Doi: 10.1172/JCI129191.

[58] Parker, W.B., 2009. Enzymology of Purine and Pyrimidine Antimetabolites Used in the Treatment of Cancer. Chemical Reviews 109(7): 2880–93, Doi: 10.1021/cr900028p.

[59] Herfarth, H.H., 2016. Methotrexate for Inflammatory Bowel Diseases - New Developments. Digestive Diseases 34(1–2): 140–6, Doi: 10.1159/000443129.

[60] Herfarth, H.H., Long, M.D., Isaacs, K.L., 2012. Methotrexate: Underused and Ignored? Digestive Diseases 30(Suppl. 3): 112–8, Doi: 10.1159/000342735.

[61] Lopez-Olivo, M.A., Siddhanamatha, H.R., Shea, B., Tugwell, P., Wells, G.A., Suarez-Almazor, M.E., 2014. Methotrexate for treating rheumatoid arthritis. Cochrane Database of Systematic Reviews, Doi: 10.1002/14651858.CD000957.pub2.

[62] Tsang-A-Sjoe, M., Bultink, I., 2015. Systemic lupus erythematosus: review of synthetic drugs. Expert Opinion on Pharmacotherapy 16(18): 2793–806, Doi: 10.1517/14656566.2015.1101448.

[63] Wagner, M., Earley, A.K., Webster, A.C., Schmid, C.H., Balk, E.M., Uhlig, K., 2015. Mycophenolic acid versus azathioprine as primary immunosuppression for kidney transplant recipients. Cochrane Database of Systematic Reviews, Doi: 10.1002/14651858.CD007746.pub2.

[64] Shaw, L.M., Korecka, M., DeNofrio, D., Brayman, K.L., 2001. Pharmacokinetic, pharmacodynamic, and outcome investigations as the basis for mycophenolic acid therapeutic drug monitoring in renal and heart transplant patients. Clinical Biochemistry 34(1): 17–22, Doi: 10.1016/S0009-9120(00)00184-3.

[65] Srinivas, T.R., Kaplan, B., Meier-Kriesche, H.-U., 2003. Mycophenolate mofetil in solid-organ transplantation. Expert Opinion on Pharmacotherapy 4(12): 2325–45, Doi: 10.1517/14656566.4.12.2325.

