## Supplemental Figures for "Systemic inhibition of *de novo* purine biosynthesis prevents weight gain and improves metabolic health by increasing thermogenesis and decreasing food intake"

**Figure S1**

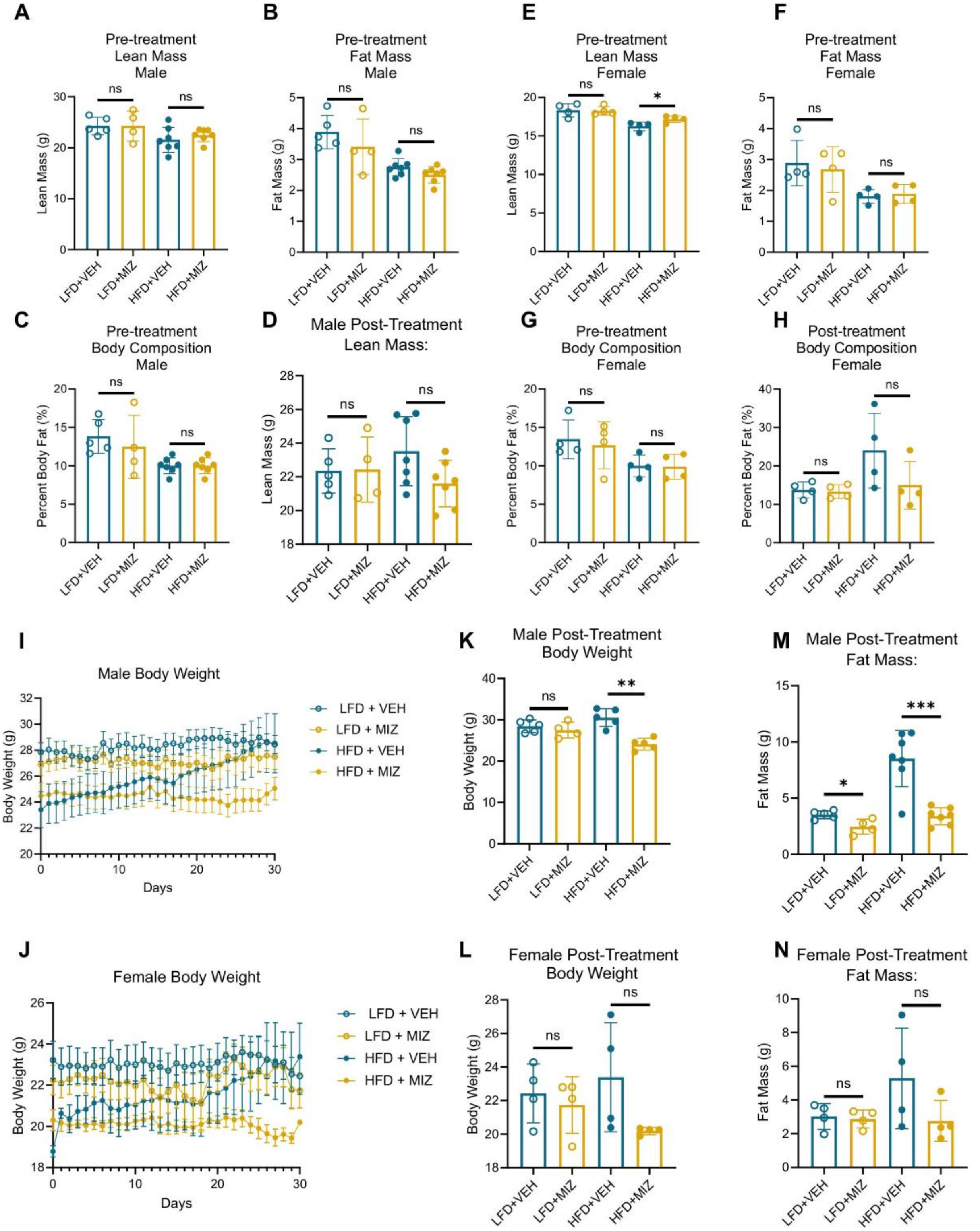


**Figure S1. Additional Body Composition Analysis.** (A – C) Quantifications of day 0 dual-energy X-ray absorptiometry (DXA) scans in male mice: lean mass (A), fat mass

(B) and body fat percentage (C). (D) Quantifications of post-treatment lean mass from dual-energy X-ray absorptiometry (DXA) scans of male mice. N = 4-7 per group. (E – G) Quantifications of day 0 dual-energy X-ray absorptiometry (DXA) scans in female mice, lean mass (E), fat mass (F) and body fat percentage (G). (H) Quantifications of post-treatment lean mass from dual-energy X-ray absorptiometry (DXA) scans of female mice. N = 4 per group. Error bars indicate mean ± standard deviation. Significance indicative of two-tailed student's *t*-test. (I – J) Absolute body weight over 30 days in male (I) or female (J) mice fed a high fat diet (HFD) or low fat diet (LFD) and treated with daily injections of mizoribine (MIZ) or vehicle (VEH). Error bars represent mean ± standard error of the mean. N = 4-5 per group. (K – L) Absolute body mass in male (K) and female (L) mice after 30 days of treatment with vehicle or mizoribine and a low or high fat diet. N = 4-5 per group. Error bars indicate mean ± standard deviation. Significance indicative of two-tailed student's *t*-test. (M – N) Quantifications of post-treatment fat mass from dual-energy X-ray absorptiometry (DXA) scans of mice in males (M) and females (N). N = 4-7 per group. Error bars indicate mean ± standard deviation. Significance indicative of two-tailed student's *t*-test. ns > 0.05, $* p \leq 0.05$, $** p \leq 0.01$, $*** p \leq 0.001$, and $**** p \leq 0.0001$.

**Figure S2**

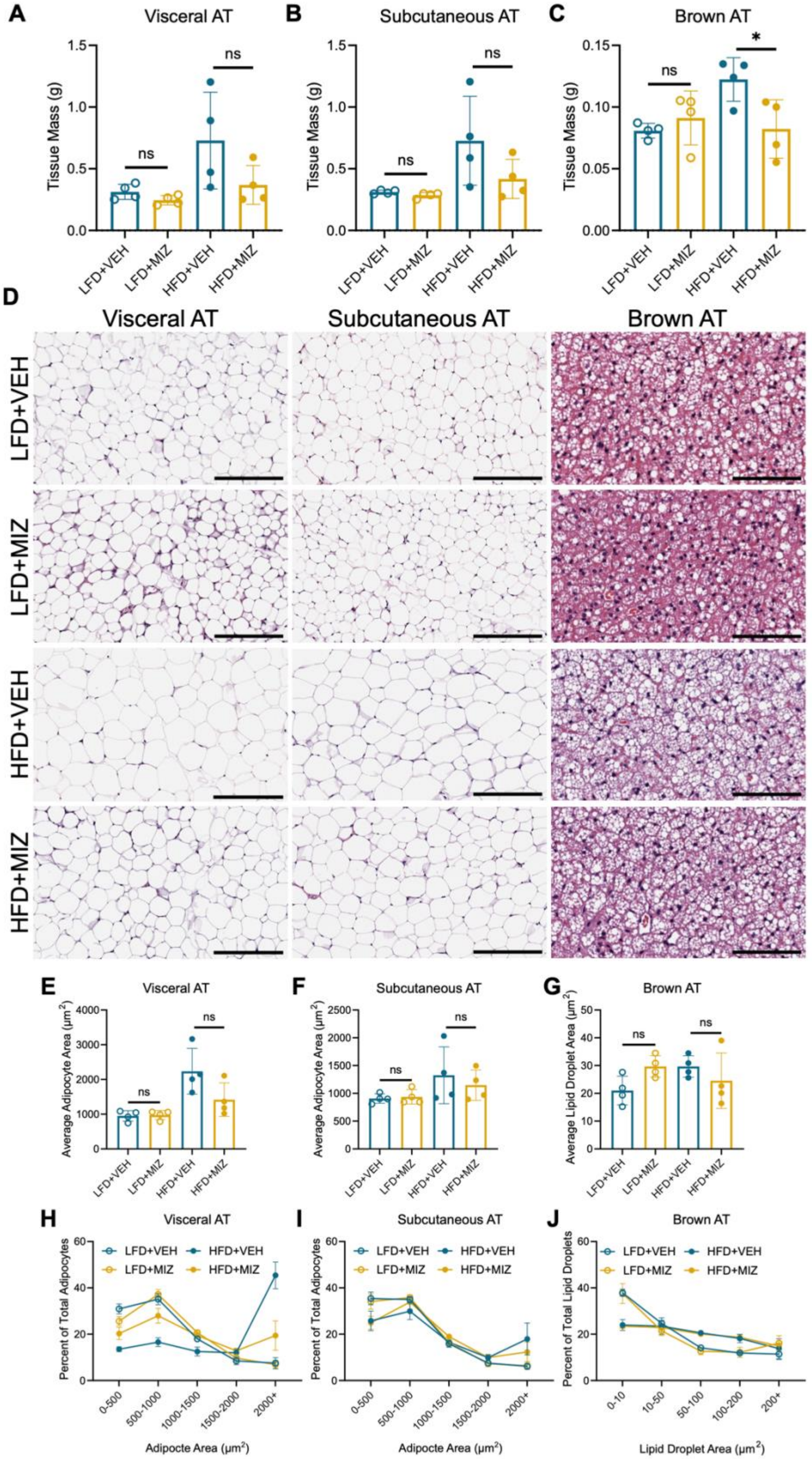
A
Visceral AT
Tissue Mass (g)
B
Subcutaneous AT
C
Brown AT
ns
*
LFD+VEH
LFD+MIZ
HFD+VEH
HFD+MIZ
D
Visceral AT
Subcutaneous AT
Brown AT
E
Average Adipocyte Area (μm²)
F
G
Average Lipid Droplet Area (μm²)
H
Percent of Total Adipocytes
Adipocte Area (μm²)
I
J
Percent of Total Lipid Droplets
Lipid Droplet Area (μm²)

**Figure S2. Terminal Adipose Tissue Masses and Morphologies from Female Mice.** (A-C) Terminal tissue mass of visceral adipose tissue (A, VAT), subcutaneous adipose tissue (B, SAT), and brown adipose tissue (C, BAT) from female mice treated for 30 days with a low fat diet (LFD) or high fat diet (HFD) and daily injections of mizoribine (MIZ) or vehicle (VEH). N = 4 per group. Error bars represent mean ± standard deviation. Significance indicative of two-tailed student's *t*-test. (D) Representative images of hematoxylin and eosin (H&E) stained sections of VAT, SAT, and BAT from female mice treated with a LFD or HFD and MIZ or VEH injections for 30 days. Scale bars = 200 µm for VAT and SAT and scale bars = 100 µm for BAT. (E-G) Quantification of visceral adipocyte (E), subcutaneous adipocyte (F), and brown adipose lipid droplet (G) size in female mice treated for 30 days with a LFD or HFD and MIZ or VEH. N = 4 per group. Error bars represent mean ± standard deviation. Significance indicative of two-tailed student's *t*-test. (H – J) Frequency distribution of visceral adipocyte (H), subcutaneous adipocyte (I), and brown adipose lipid droplet (J) size in female mice treated for 30 days with a LFD or HFD and MIZ or VEH. N = 4-5 per group. ns > 0.05, ∗ $p \leq 0.05$, ∗∗ $p \leq 0.01$, ∗∗∗ $p \leq 0.001$, and ∗∗∗∗ $p \leq 0.0001$.

**Figure S3**

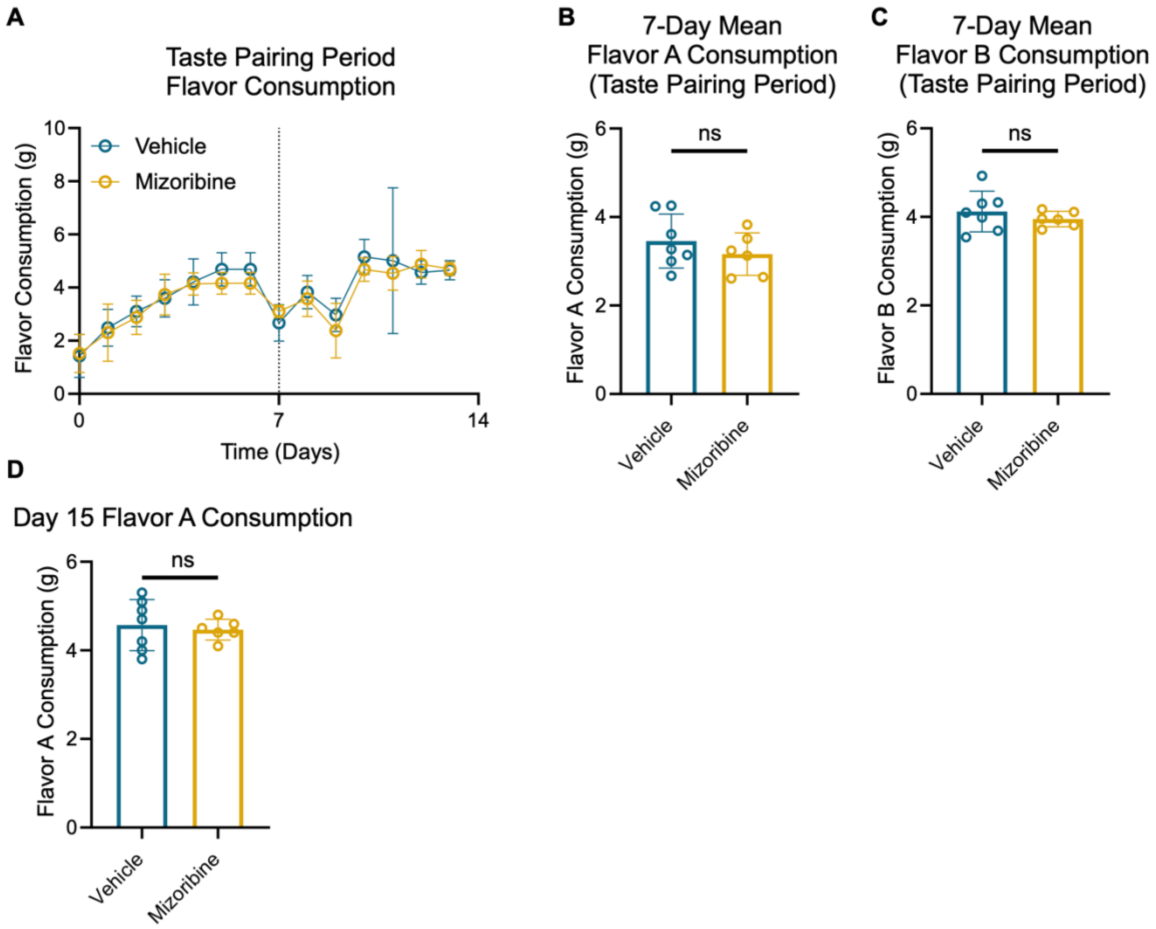


**Figure S3. Conditioned Taste Aversion Controls.** (A) Daily flavor consumption during the control and experimental taste pairing periods. (B – C) Mean daily consumption during the control (B) and experimental (C) taste pairing periods. (D) Measurement of flavor A consumption on day 15 of the experiment, comparing conditioned taste aversion to flavor A associated with mizoribine versus vehicle treatment. Error bars represent mean ± standard deviation. Significance indicative of two-tailed student's *t*-test. ns > 0.05, $* p \leq 0.05$, $** p \leq 0.01$, $*** p \leq 0.001$, and $**** p \leq 0.0001$.

**Figure S4**

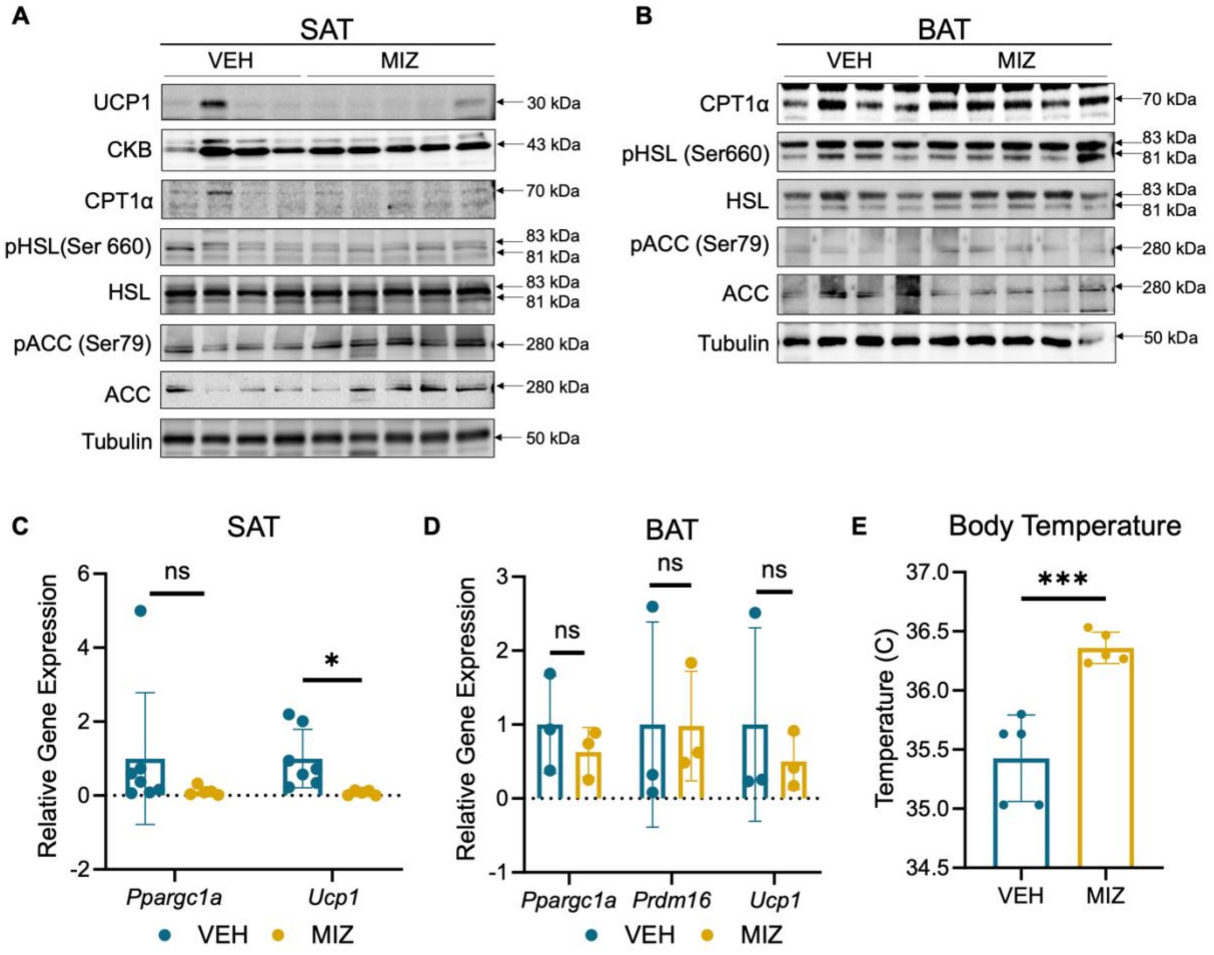


**Figure S4: Brown and Subcutaneous White Adipose Tissue Phenotype in Mizoribine-Treated Mice.** (A) Immunoblot analysis of UCP1, CKB, CPT1α, p-HSL (Ser660), HSL, p-ACC (Ser79), ACC, and Tubulin in subcutaneous adipose tissue from male mice fed a high fat diet and treated with mizoribine or vehicle injections. (B) Immunoblot analysis of CPT1α, p-HSL (Ser660), HSL, p-ACC (Ser79), ACC, and Tubulin in brown adipose tissue from male mice fed a high fat diet and treated with mizoribine or vehicle injections. (C – D) Gene expression analysis of *Ppargc1a*, *Prdm16*, and *Ucp1* in subcutaneous (C) and brown (D) adipose tissue (normalized to 16S rRNA). N = 3-7 per group. Error bars represent mean ± standard deviation. Significance indicative of two-tailed student's *t*-test. (E) Body temperature in mice after treatment with a high fat diet and mizoribine or vehicle for 30 days (measured by implanted temperature probes). Measurements representative of 3-day average body temperature. N = 5. Error bars represent mean ± standard deviation. Significance indicative of two-tailed student's *t*-test. ns > 0.05, $* \ p \leq 0.05$, $** \ p \leq 0.01$, $*** \ p \leq 0.001$, and $**** \ p \leq 0.0001$.

**Figure S5**

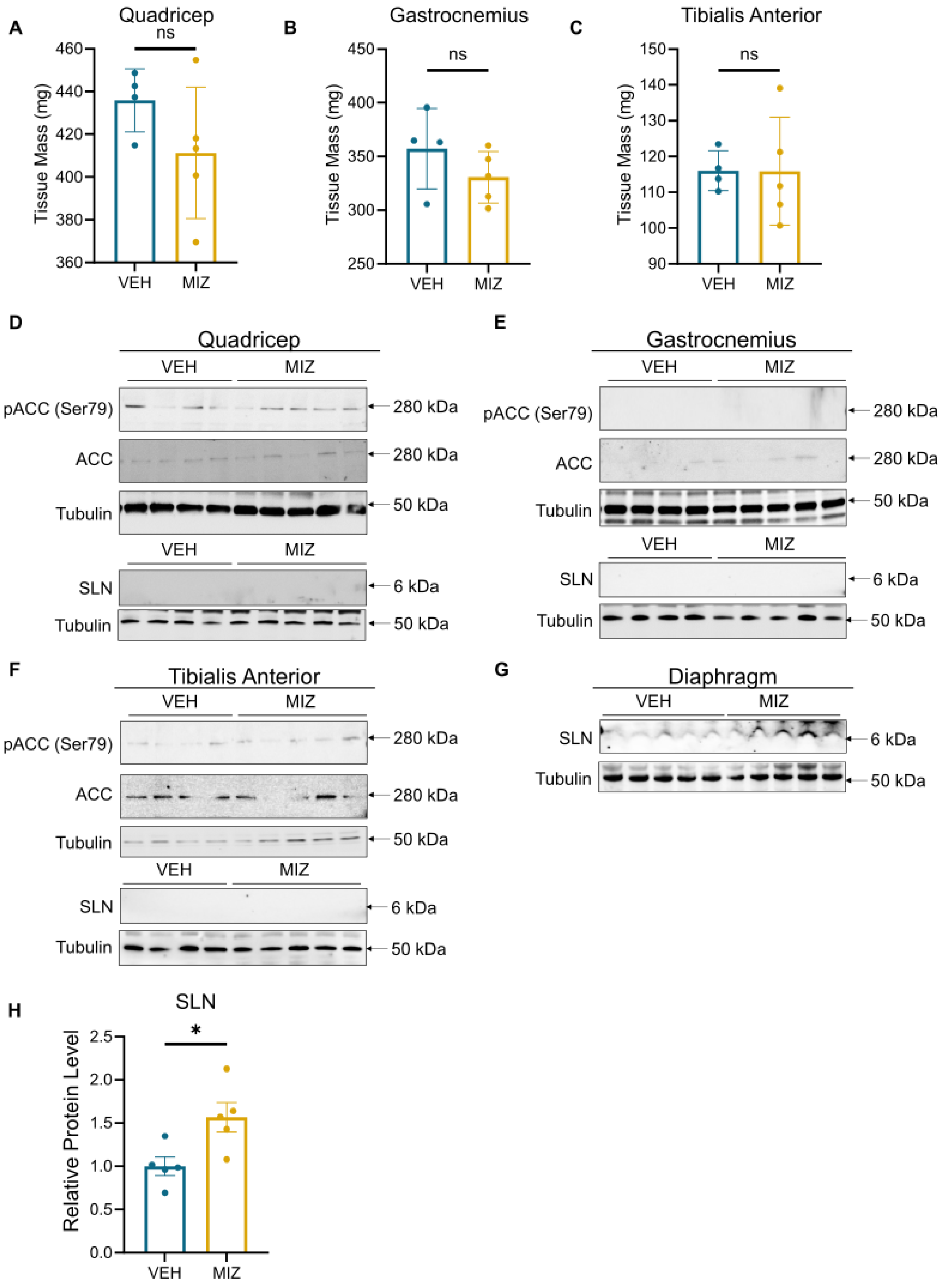
A
Quadricep
ns
Tissue Mass (mg)
460
440
420
400
380
360
VEH
MIZ
B
Gastrocnemius
ns
Tissue Mass (mg)
450
400
350
300
250
VEH
MIZ
C
Tibialis Anterior
ns
Tissue Mass (mg)
150
140
130
120
110
100
90
VEH
MIZ
D
Quadricep
VEH
MIZ
pACC (Ser79)
280 kDa
ACC
280 kDa
Tubulin
50 kDa
VEH
MIZ
SLN
6 kDa
Tubulin
50 kDa
E
Gastrocnemius
VEH
MIZ
pACC (Ser79)
280 kDa
ACC
280 kDa
Tubulin
50 kDa
VEH
MIZ
SLN
6 kDa
Tubulin
50 kDa
F
Tibialis Anterior
VEH
MIZ
pACC (Ser79)
280 kDa
ACC
280 kDa
Tubulin
50 kDa
VEH
MIZ
SLN
6 kDa
Tubulin
50 kDa
G
Diaphragm
VEH
MIZ
SLN
6 kDa
Tubulin
50 kDa
H
SLN
*
Relative Protein Level
2.5
2.0
1.5
1.0
0.5
0.0
VEH
MIZ

**Figure S5: Muscle Phenotype in Mizoribine-Treated Mice.** (A – C) Muscle mass of quadricep (A), gastrocnemius (B), tibialis anterior (C), and from male mice fed a high fat diet and treated with injections of vehicle or mizoribine. N = 4-5. Error bars represent mean ± standard deviation. Significance indicative of two-tailed student's *t*-test. (D – F) Immunoblot analysis of SLN, p-ACC (Ser79), ACC, and Tubulin in the quadricep (D), gastrocnemius (E), and tibialis anterior (F) from male mice fed a high fat diet and treated with injections of vehicle or mizoribine. (G) Immunoblot analysis of SLN and Tubulin in the diaphragm of male mice fed a high fat diet and treated with injections of vehicle or mizoribine. (H) Quantification of SLN (normalized to Tubulin) in the diaphragm of male mice fed a high fat diet and treated with injections of vehicle or mizoribine N = 5. Error bars represent mean ± standard deviation. Significance indicative of two-tailed student's *t*-test. ns > 0.05, $*$ $p \leq 0.05$, $**$ $p \leq 0.01$, $***$ $p \leq 0.001$, and $****$ $p \leq 0.0001$.

**Figure S6**

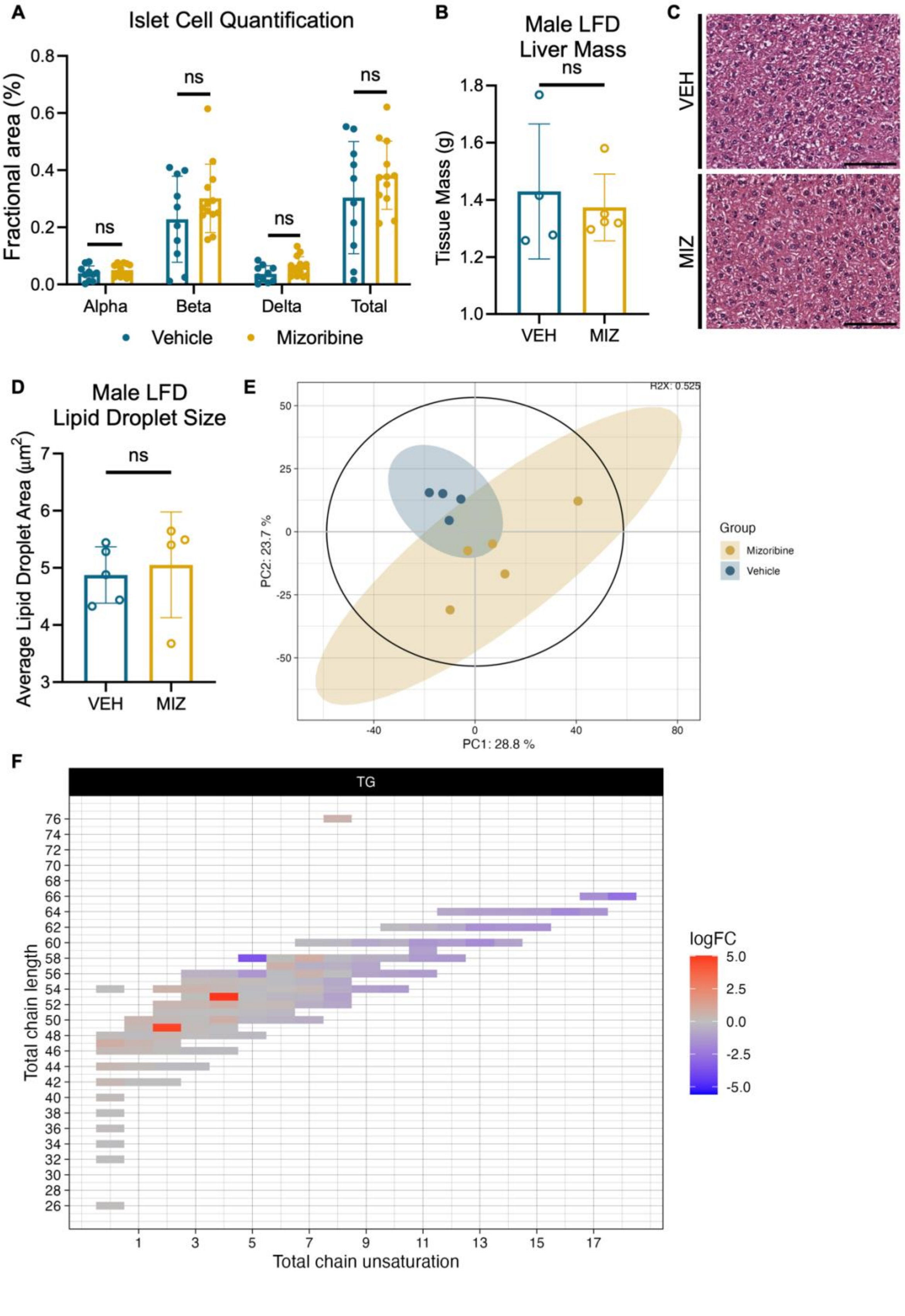
A
Islet Cell Quantification
Fractional area (%)
0.8
0.6
0.4
0.2
0.0
ns
Alpha
Beta
Delta
Total
Vehicle
Mizoribine
B
Male LFD
Liver Mass
Tissue Mass (g)
1.8
1.6
1.4
1.2
1.0
VEH
MIZ
C
VEH
MIZ
D
Male LFD
Lipid Droplet Size
Average Lipid Droplet Area (μm²)
7
6
5
4
3
E
R2X: 0.525
PC1: 28.8 %
PC2: 23.7 %
Group
Mizoribine
Vehicle
F
TG
Total chain length
Total chain unsaturation
logFC
5.0
2.5
0.0
-2.5
-5.0

**Figure S6: Effects of Mizoribine Treatment on Hepatic and Circulating Lipids.** (A) Quantification of immunofluorescent labeled pancreatic islet endocrine cells. N = 10-13. Error bars represent mean ± standard deviation. Significance indicative of two-tailed student's *t*-test. (B) Terminal liver mass from male mice treated with a low fat diet (LFD) and daily injections of mizoribine (MIZ) or vehicle (VEH) for 30 days. (C) Representative images of hematoxylin and eosin (H&E) stained sections of liver from male mice treated with a LFD and either vehicle or mizoribine injections. (D) Quantification of hepatic lipid droplet size in male mice treated with a LFD and daily injections of mizoribine (MIZ) or vehicle (VEH) for 30 days. N = 4-5 per group. Error bars represent mean ± standard deviation. Significance indicative of two-tailed student's *t*-test. (E) Principal component analysis of the plasma lipidome of male mice treated with vehicle or mizoribine and a HFD. (F) Heat map of differential plasma triglycerides abundance by chain length and saturation. Heat map color depicts $log_2$FoldChange where a positive $log_2$FoldChange indicates increased abundance in mizoribine samples and a negative $log_2$FoldChange indicates decreased abundance in mizoribine samples relative to vehicle samples. Lipidomic data from male mice treated with vehicle or mizoribine and a HFD. ns > 0.05, $* \ p \leq 0.05$, $** \ p \leq 0.01$, $*** \ p \leq 0.001$, and $**** \ p \leq 0.0001$.
